# Catabolite Activator Protein and quorum sensing cross-control group behaviors in *Vibrio campbellii*

**DOI:** 10.64898/2026.07.29.741505

**Authors:** Chase M. Mullins, Alyssa S. Ball, Logan J. Geyman, Lydia K. Lukich, Lydia K. Hermann, Ram Podicheti, Zhongqing Ren, Xindan Wang, Douglas B. Rusch, Julia C. van Kessel

## Abstract

*Vibrio* species adapt to different niches by sensing and responding to environmental signals such as nutrients, host cues, and quorum sensing autoinducers. *Vibrio campbellii* uses the master quorum sensing transcription factor LuxR to control expression of hundreds of genes at high cell density. Furthermore, many gamma-proteobacteria use the global transcription factor Catabolite Activator Protein (CAP) to control numerous physiologically relevant pathways, many of which are also modulated by quorum sensing, such as competence, biofilm formation, and carbon metabolism. However, the extent to which these two global transcription factors overlap to co-regulate gene expression and bacterial behaviors is understudied. In this work, we used ChIP-seq and RNA-seq to determine the individual and combined regulons of CAP and LuxR in *V. campbellii*. We found that CAP and LuxR co-occupied 11 promoters to synergistically or antagonistically co-regulate genes involved with metabolism, respiration, and virulence. It was previously proposed that CAP and LuxR both bound the bioluminescence (*luxCDABE*) promoter to co-regulate these genes, and our RNA-seq data showed that these were indeed the most strongly co-regulated genes. However, our ChIP-seq data revealed that only LuxR bound the *luxCDABE* promoter *in vivo*. This pattern of co-regulation—where both CAP and LuxR strongly impact transcription but only LuxR binds the promoter—was the most common mechanism observed. Our model for bioluminescence regulation is that CAP indirectly activates *luxCDABE* expression through the regulation of an intermediate factor. This study established new connections between the global gene regulatory networks underpinning nutrient sensing and population sensing.

**Importance:** *Vibrio* bacteria (vibrios) colonize and infect diverse marine hosts including corals, fish, oysters, and shrimp, and can also cause life-threatening human infections, all of which are rising annually due to increasing ocean temperatures. To develop effective treatments against *Vibrio* infections, it is critical to understand the global gene regulation mechanisms vibrios use that enable pathogenic lifestyles. Signal transduction systems in bacteria are well-characterized; however, how vibrios coordinate global gene expression changes in response to multiple environmental inputs is understudied. Here, we determined how the aquaculture pathogen *Vibrio campbellii* regulates global gene expression in response to bacterial population signals and nutrient availability signals. Our findings help contextualize how vibrios respond to their fluctuating environments in nature.

## Introduction

*Vibrio* bacteria (vibrios) are versatile organisms that exist in free-living planktonic states, form complex microbial communities called biofilms, and colonize and infect the guts of diverse hosts, including oysters, fish, shrimp, corals, and humans (1–6). *Vibrio* infections of animals in native marine ecosystems and artificial farms cause devastating consequences for aquaculture industries, coral reefs, and human health (3–7). Vibrios, like many bacteria, are effective pathogens because of their ability to quickly and efficiently sense and respond to environmental signals, like nutrient availability, host signals, and quorum sensing autoinducers (8, 9). As *Vibrio* bacteria move between lifestyles, they employ numerous gene regulation mechanisms to optimize nutrient utilization and change behaviors (1, 2, 8, 10–12). Most notably, vibrios use the population-density signaling system called quorum sensing (QS) to coordinate these lifestyle switches and activate pathogenic behaviors (8, 11, 12). Indeed, *Vibrio* species have emerged as models for studying QS gene regulation due to their well-defined QS signaling cascades and genetic tractability (10, 12). However, how QS intersects with other environmental signaling systems, like nutrient sensing systems, within the *Vibrio* genus is an understudied topic.

QS depends on the production and detection of small signaling molecules called autoinducers (AIs), which accumulate in the environment as a function of cell density and enable cells to detect the number and type of bacterial cells nearby (11). In vibrios, AI concentrations ultimately control expression of the TetR-like master transcriptional regulator called LuxR/HapR, conserved in all studied *Vibrio* species (10). Here, we focus on the QS system in the model shrimp pathogen *Vibrio campbellii* BB120 (also known as ATCC BAA-1116 and previously classified as *Vibrio harveyi*) (13). The LuxR protein in *V. campbellii* is distinct from LuxR in *Aliivibrio fischeri* and other bacteria, which regulates gene expression upon binding acyl-homoserine lactone autoinducers produced from the autoinducer synthase LuxI (11, 12). In *V. campbellii*, at low cell densities (low AIs), LuxR levels are low and control ∼80 genes. Conversely, at high cell densities (high AIs), LuxR increases ∼10-fold and regulates >400 genes, including those required for virulence and bioluminescence (10). Numbers of *Vibrio* bacteria in planktonic environments vary depending on environmental conditions, like temperature and salinity (3, 14, 15). However, *Vibrio* bacteria frequently reach high cell concentrations when growing in host organisms, like in oysters, bobtail squid, or shellfish, which provide sources of carbon, nitrogen, fats, and the opportunity to scavenge iron, all of which are scarce in planktonic environments (5, 15, 16). Thus, it is important to understand how vibrios integrate both population density signals and nutrient availability signals to enable the colonization, proliferation, and infection of hosts.

Bacteria possess numerous and diverse mechanisms for directly and indirectly sensing nutrient availability. In this study, we focus on carbon availability sensing via the Catabolite Activator Protein (CAP) which is also known as the cyclic-AMP receptor protein (CRP). CAP has been historically studied in *Escherichia coli* as a transcription factor that binds cyclic-AMP to regulate alternative carbon source utilization (*e.g*., lactose, maltose, galactose) when preferred carbon sources like glucose are limiting (17–24). Numerous studies have broadened our understanding of CAP beyond its historical role and revealed that CAP is a global regulator in many bacterial pathogens (9). For example, CAP regulates virulence factors, QS genes, and carbon utilization genes in *Yersinia pestis*, *Mycobacterium tuberculosis,* and *Vibrio cholerae* (9, 25–34). Specifically in *V. cholerae*, CAP regulates several QS genes including *hapR* and *cqsA,* the master QS transcription factor and the CAI-1 autoinducer synthase, respectively (30, 35). Additional studies confirmed that CAP-like proteins are deeply intertwined with QS, like how the global regulator Vfr in the pathogen *Pseudomonas aeruginosa* controls expression of both the *las* and *rhl* QS systems (36).

Several studies in different *Vibrio* species have demonstrated that CAP and LuxR homologs can directly overlap at the promoters for QS genes, metabolism genes, and virulence genes. For instance, in *V. cholerae,* the LuxR homolog HapR overlaps with CAP at seven sites across the chromosome, including at the *murPQ* promoter, where CAP and HapR repress genes important for metabolizing chitinous surfaces in the environment (37, 38). Another study in *V. cholerae* found that HapR and CAP bind to the *prtV* metalloprotease promoter to regulate an intragenic small RNA that is important for bacteriophage CTXΦ activation and cell lysis (39). Furthermore, in *Vibrio vulnificus*, the LuxR homolog SmcR overlaps with CAP at the *vvpM* and *vvpE* promoters to repress metalloprotease production and activate elastase production, respectively, which are both important for infection (40–42). Lastly, in *V. campbellii*, it was proposed that CAP and LuxR both bind to the *luxCDABE* promoter to regulate bioluminescence at high cell densities (43). These studies highlight that CAP and LuxR homologs can overlap at specific promoters to co-regulate gene expression in closely related *Vibrio* species. However, it is not clear if CAP and LuxR co-regulation is a conserved, global mechanism employed by *Vibrio* species, and whether this regulation controls specific pathways.

To address this question, we defined the global co-regulon between CAP and LuxR in the model QS organism *V. campbellii* using RNA-sequencing and Chromatin Immunoprecipitation-sequencing. We discovered that the co-regulated genes with the largest fold-changes by both LuxR and CAP were those encoding for bioluminescence (*luxCDABE; lux*), which is a native phenotype and historical model for studying QS-gene regulation. We provide evidence to refute a previously proposed model where CAP and LuxR both bind the *lux* promoter to activate expression and instead propose our own model where LuxR binds the *lux* promoter, and CAP regulates expression of an intermediate regulator required for *lux* expression. Beyond the *luxCDABE* locus, we discovered that CAP and LuxR overlap at a dozen promoters and utilize several mechanisms to co-regulate global gene expression. Ultimately, our research supports the hypothesis that vibrios use CAP and LuxR/HapR to control global gene expression in response to fluctuations in nutritional and populational cues, respectively.

## Results

### CAP and LuxR both regulate hundreds of genes

Based on the findings from the Walker *et al.* study that demonstrated an important role for CAP in co-regulation with HapR in *V. cholerae* (37), we hypothesized that CAP also plays a co-regulatory role with LuxR in *V. campbellii*. To identify the global transcriptional overlap between CAP and LuxR, we performed RNA-seq in wild-type (WT), Δ*cap*, Δ*luxR*, and Δ*cap* Δ*luxR V. campbellii* BB120 strains on RNA collected at high cell density (OD_600_ = 1.0). For all our comparisons, we defined significant hits as those genes that were differentially regulated ≥4-fold (false discovery rate [FDR] = 0.05, *p*-value < 0.05). We first defined the CAP regulon by comparing transcripts between Δ*cap* and WT and discovered that CAP regulated 669 genes: 468 were activated and 201 were repressed (Figs. 1A, 1E, Dataset S1). The *V. campbellii* CAP regulon contained expected genes like those involved in maltose utilization and adenylate cyclase, similar to the CAP regulon in *E. coli* (44). We validated the results from our RNA-seq using RT-qPCR and demonstrated that the observed gene expression differences were complemented by expressing *cap* in the Δ*cap* strain (Fig. S1). When we compared the gene expression profiles of Δ*luxR* to WT, we observed similar results to previously published RNA-seq and microarray datasets of the LuxR regulon (45–47). The results in this dataset showed that LuxR regulated 266 genes: 121 were activated and 145 were repressed (Figs. 1B, 1E, Dataset S1). The overlap between CAP and LuxR regulons consisted of 97 genes, and the most highly co-regulated genes were those required for bioluminescence, *luxCDABE* (Figs. 1A, 1B, 1E, Dataset S1). CAP and LuxR were also found to co-regulate genes that encode transcription factors and metabolite/sugar transporters, as well as genes implicated in processes like polysaccharide synthesis, respiration, and metabolism (Dataset S1).

**Figure 1.**
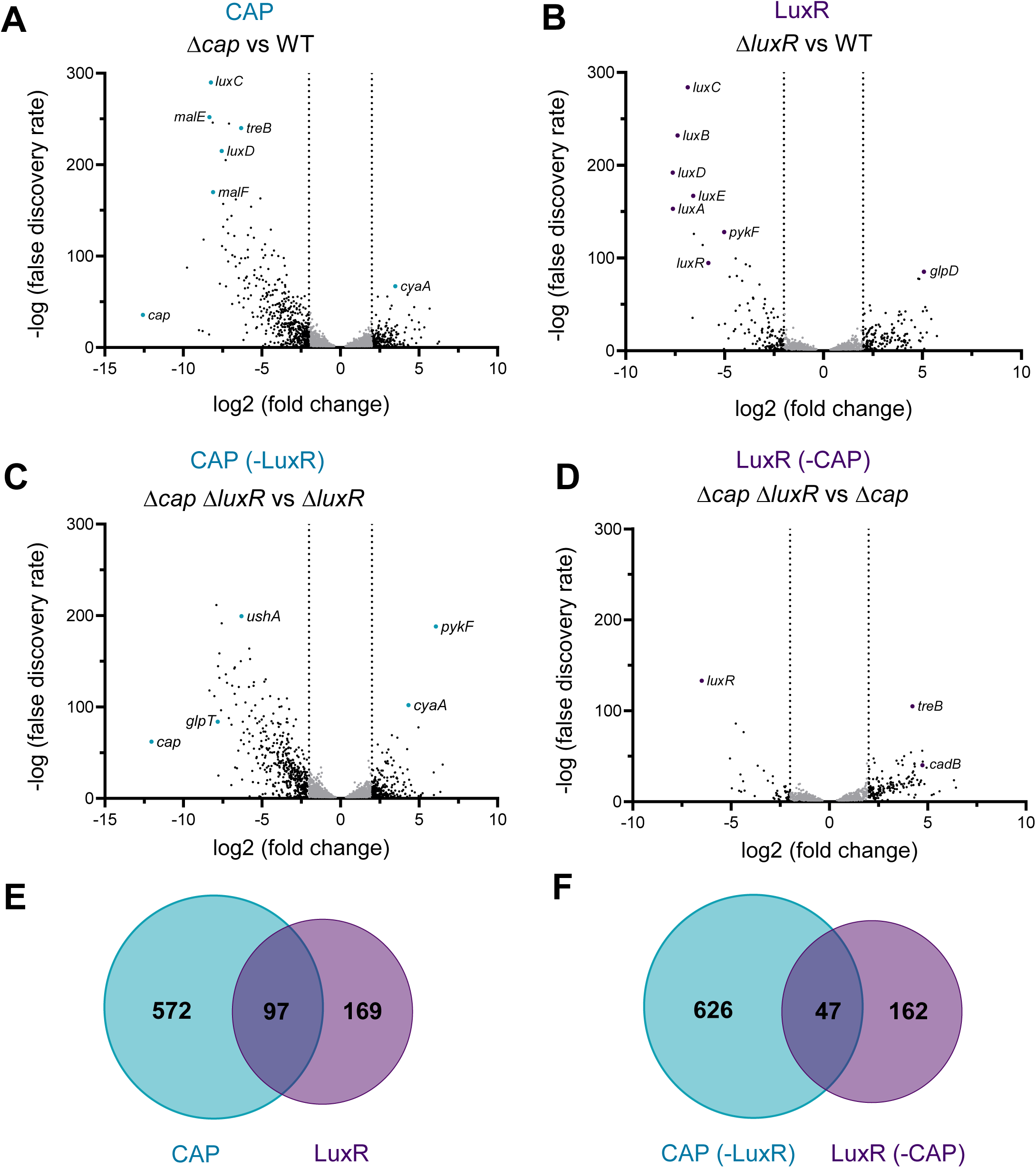
CAP and LuxR co-regulate genes in *V. campbellii*. (A-D) RNA-seq was performed on RNA collected from WT, Δ*cap,* Δ*luxR, and* Δ*cap* Δ*luxR V. campbellii strains* grown to high cell density (OD_600_ of 1.0). Volcano plots showing the *q*-value (-log BH [Benjamini-Hochberg] adjusted *p-*value) over the log2-fold change for Δ*cap* vs WT (A), Δ*luxR* vs WT (B), Δ*cap* Δ*luxR* vs Δ*luxR* (C), and Δ*cap* Δl*uxR* vs Δ*cap* (D) are shown. Select genes of interest are labeled in each graph. Each point on the graph is the average from 3 biological replicates. A negative log2-fold value indicates genes that are activated, and a positive log2-fold value indicates genes that are repressed. Note: the *q*-value for *luxC* in the Δ*cap* strain could not be calculated because the BH adjusted *p*-value was calculated to be 0. For graphing purposes, we set the value to be the same as *luxC* from the Δ*luxR* vs WT comparison. (E-F) Venn diagrams showing the overlap of significantly (≥4-fold) regulated genes between CAP and LuxR (E) and CAP (-LuxR) and LuxR (-CAP) (F). The CAP regulon is represented in teal and LuxR in purple.

Next, we assessed each protein’s individual contribution to the transcriptome by comparing the Δ*cap* Δ*luxR* strain to either the Δ*cap* or the Δ*luxR* strains. The CAP regulon in the absence of LuxR (673 genes) comprised 457 activated genes and 216 repressed genes (Figs. 1C, 1F, Dataset S1). The LuxR regulon in the absence of CAP (209 genes) comprised 49 activated genes and 160 repressed genes (Figs. 1D, 1F, Dataset S1). The overlap between the individual regulons of CAP and LuxR consisted of 47 genes, including those encoding for predicted acetyltransferases, transcription factors, bacterioferritin, trehalose utilization, and fumarate reductase (Fig. 1F, Dataset S1). Comparing the two co-regulon datasets, and accounting for shared genes, we determined that CAP and LuxR co-regulated 136 total genes in *V. campbellii.* Collectively, these RNA-seq data 1) established a LuxR and CAP co-regulon, 2) determined each protein’s individual influence over the transcriptome, and 3) identified that the most highly regulated genes by both CAP and LuxR were *luxCDABE*.

### CAP and LuxR are both required for bioluminescence production

Our RNA-seq data implicated the *luxCDABE* operon as the strongest co-regulated genes in *V. campbellii.* The *V. campbellii luxCDABE* promoter (P*_lux_*) has been extensively studied over the past forty years (43, 45, 47–54). The Chatterjee *et al.* study also identified CAP and LuxR as co-activators of P*_lux_* (43). However, the mechanism of co-regulation of this key operon by both LuxR and CAP was untested. We measured bioluminescence in WT, Δ*cap*, and Δ*luxR* strains throughout a growth curve to determine the effect of each regulator on bioluminescence production over time. For the WT *V. campbellii* strain, bioluminescence production increased as OD_600_ increased (Fig. 2A), as historically shown (11, 50, 55–59). Once autoinducer concentrations reached quorum (OD_600_ = ∼0.2-0.4 as measured on a plate reader, which corresponds to ∼1.0 in a standard spectrophotometer), bioluminescence increased accordingly due to LuxR-dependent activation of the *lux* operon (10, 47, 49). To illustrate this point, a Δ*luxR* strain was incapable of producing bioluminescence which corresponded to a >1000-fold decrease in *luxC* transcript levels, both of which were restored to WT levels by complementation of *luxR* on an episomal plasmid (Figs. 2A, 2B). The Δ*cap* strain also exhibited no bioluminescence production and >100-fold lower *luxC* transcript levels, and both were restored to WT levels after complementation (Figs. 2A, 2B). When either *cap* or *luxR* expression was induced in a WT background, bioluminescence increased earlier in the time-course than WT levels, confirming that both proteins are activators (Fig. 2A). Neither CAP nor LuxR were sufficient for bioluminescence production, even under over-expression conditions (Figs. 2A, 2B).

**Figure 2.**
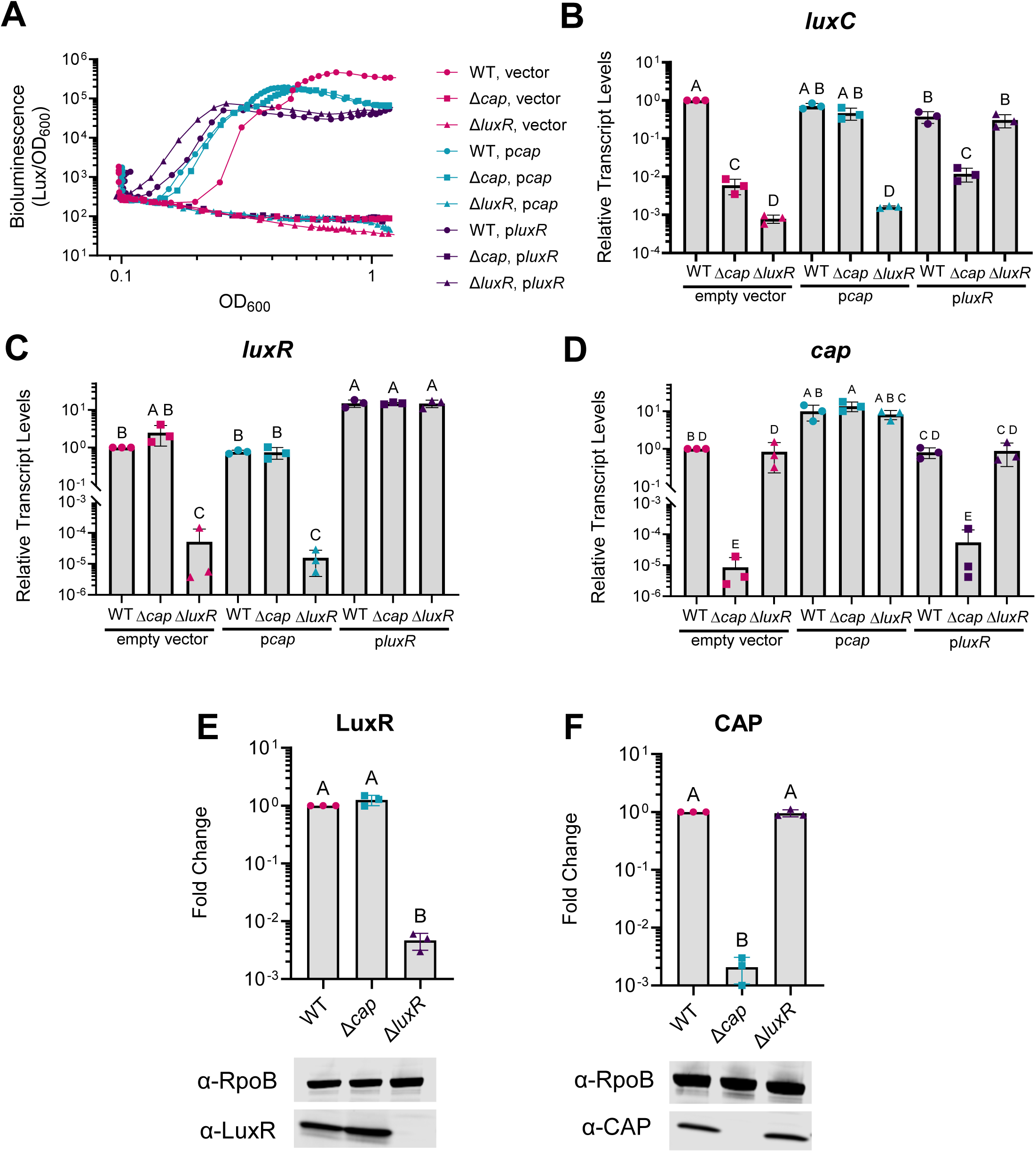
CAP and LuxR are both required for bioluminescence production. (A) Bioluminescence production and OD_600_ were measured for 24 hours for Wild-type (WT), Δ*luxR*, and Δ*cap V. campbellii* strains harboring either an empty vector (pCS027) or an episomal plasmid expressing *cap* (pAB100) or *luxR* (pJV388) from an IPTG inducible promoter (25 µM). The figure is a representative graph from three biological replicates and each line on the graph is an average of two technical replicates. The other biological replicates are shown in supplemental figure 5. (B, C, D) RT-qPCR analysis of RNA collected at an OD_600_ of 1.0 from the same strains and conditions as panel A. Relative transcript levels were measured for *luxC* (B), *luxR* (C), and *cap* (D) using the ΔΔC_T_ method and *hfq* as an internal control. (E, F) Western blot analysis of lysates using LuxR polyclonal antibodies (α-LuxR; E) or CAP monoclonal antibodies (α-CAP; F). RNA polymerase subunit B (α-RpoB) monoclonal antibodies were used to detect RpoB as a loading control. Fold change was calculated by normalizing CAP and LuxR to their respective RpoB signals and then comparing to the WT sample in each group. Three biological replicates are represented on the graphs, and the qualitative images are a representative from one biological replicate. Full membrane images from all three biological replicates are shown in supplemental figure 6. (B-F) A one-way ANOVA test was performed on lognormally distributed data (Shapiro-Wilk test), followed by Tukey’s multiple comparisons test. Different letters above bars indicate differences in significance between strains in pairwise comparisons (*p*<0.05; *n*=3). Error bars represent the standard deviation.

One possible explanation for the requirement of both LuxR and CAP as activators of *luxCDABE* was that LuxR or CAP regulated the transcription and/or protein level of the other. We measured *luxR* and *cap* transcript and protein levels in Δ*cap* and Δ*luxR* strains compared to WT. We observed no change in *luxR* transcripts or LuxR protein levels in the absence of CAP (Fig. 2C, 2E). Additionally, *cap* transcripts and protein levels were similar between WT and Δ*luxR* strains (Fig. 2D, 2F). From these collective data, we concluded that 1) CAP and LuxR are both necessary but not sufficient for *lux* operon transcription and bioluminescence production, and 2) CAP and LuxR do not control expression of each other under the conditions we tested.

Another possible mechanism for CAP and LuxR to affect bioluminescence production was that CAP regulated a required substrate or intermediate. The long-chain fatty aldehyde substrate that the Luciferase enzyme (LuxAB) requires to produce bioluminescence is produced from the LuxCDE reduction of fatty acid substrates, which are produced from the fatty acid synthase pathway. Additionally, the bioluminescence reaction requires reduced flavin mononucleotide to serve as a redox cofactor (60, 61). To test if CAP and LuxR affected substrate accumulation, we genetically separated the *luxCDABE* genes from their native promoter and placed their expression under the control of an IPTG-inducible promoter (P*_tac_*). We then measured light production in WT, Δ*cap*, and Δ*luxR* strains harboring our inducible bioluminescence system. We observed constitutive bioluminescence even in the absence of the inducer for all three strains (Fig. S2A), likely due to leaky expression from the P*_tac_* promoter.

Although the Δ*cap* and Δ*luxR* strains were bright upon induction, they produced 10-fold less light than the WT strain due to a decrease in bioluminescence production near stationary phase that did not occur in the WT strain (calculated using an area under the curve analysis; Fig. S2B). From these data, we concluded that the major impact of CAP and LuxR on bioluminescence is through *luxCDABE* transcriptional control rather than affecting the accumulation of substrate or metabolic intermediates.

### Bioluminescence is activated in a cyclic-AMP-dependent manner

Because CAP regulates transcription upon binding the intracellular signaling nucleotide cyclic-AMP (cAMP) which is made by adenylate cyclase (CyaA), we investigated the role of cAMP and CyaA on bioluminescence production (9, 19, 23). Based on the literature in *E. coli*, we expected that a *cyaA* mutant would phenocopy a *cap* mutant and not be bioluminescent.

Surprisingly, the Δ*cyaA* strain produced bioluminescence, although it was not as maximally bright and was significantly delayed in bioluminescence production compared to WT (Figs. 3A, 3B). However, upon chromosomal complementation, bioluminescence was restored to WT levels in the Δ*cyaA* strain (Figs. 3A, 3B).

**Figure 3.**
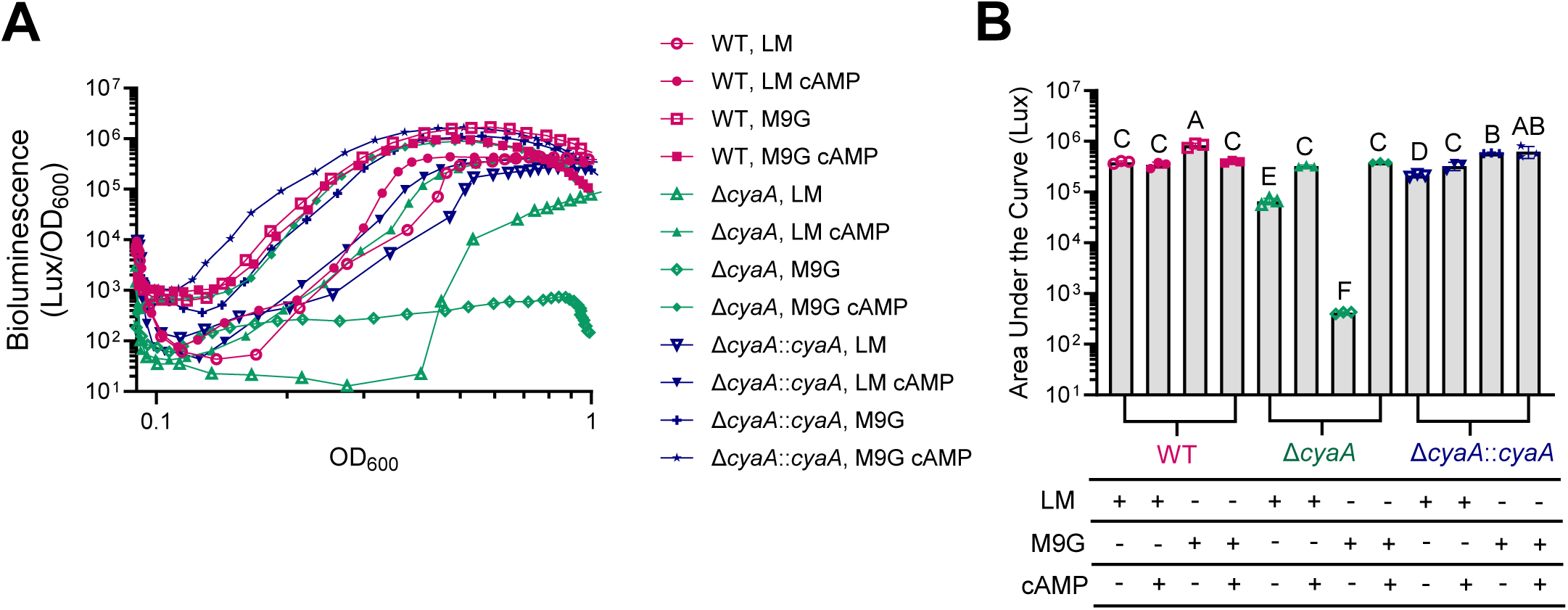
Bioluminescence production requires cyclic-AMP. (A) Bioluminescence production and OD_600_ were measured over 24 hours for WT or Δ*cyaA V. campbellii* strains grown in either a rich, undefined medium (LM; LB + NaCl) or M9 minimal medium supplemented with 20 mM glucose (M9G). The Δ*cyaA* strains were complemented with an IPTG-inducible (10 µM) copy of *cyaA* inserted on the chromosome at a non-polar site (Δ*cyaA*::*cyaA*). Exogenous cyclic-AMP (cAMP) was added to the medium at a final concentration of 1 mM. The graph is a representative of three biological replicates and each line on the graph is the average of two technical replicates. Additional biological replicates are presented in supplemental figure 7. (B) Area under the curve (Lux) was calculated from the three biological replicates in A and Fig. S7. A one-way ANOVA test was performed on normally distributed log-transformed data (D’Agostino-Pearson test on the residuals) followed by Tukey’s multiple comparisons test. Different letters above bars indicate differences in significance between strains in pairwise comparisons (*p*<0.05; *n*=3). Error bars represent the standard deviation. Plusses and minuses under the bars indicate if the strain was grown in LM or M9G with or without cAMP.

Previous studies have demonstrated that *Vibrio* species can sense and respond to extracellular cAMP, although the mechanism is unclear (35, 62–64). We exploited this property by providing extracellular cAMP in the growth medium and observing the effect on bioluminescence. Indeed, extracellular cAMP stimulated earlier bioluminescence production for a WT strain when compared to the same medium without cAMP (Fig. 3A). Providing extracellular cAMP to a Δ*cyaA* strain restored bioluminescence to WT levels, demonstrating that the dark phenotype was due to a lack of cAMP (Figs. 3A, 3B). These experiments led us to the important realization that our undefined medium, called LM for Lysogeny Broth Marine (LB with 2% extra NaCl), which contains yeast extract, may contain residual amounts of cAMP because yeast also use cAMP for signaling (62). We reasoned that if we grew Δ*cyaA* in a defined medium lacking cAMP, like M9 minimal medium supplemented with glucose (M9G), bioluminescence would be inhibited. Indeed, a Δ*cyaA* strain was dark in M9G, confirming that cAMP from the yeast extract in LM was stimulating a minor bioluminescence response (Figs. 3A, 3B). Another reason we chose to test for bioluminescence production in M9G is because glucose has been historically shown to repress bioluminescence in *Aliivibrio fischeri,* presumably through carbon catabolite repression via CAP (63, 65). However, a WT *V. campbellii* strain was not dark nor delayed in M9G; rather, it was brighter and began to bioluminesce earlier than a WT strain grown in LM (Figs. 3A, 3B). Collectively from these experiments, we concluded that bioluminescence is a cyclic-AMP-dependent phenotype and that glucose activates bioluminescence in *V. campbellii*.

### CAP is predicted to bind P*_lux_*

We demonstrated that CAP is a key activator of the *lux* operon, and previous work has shown that CAP can bind P*_lux_* DNA in electrophoretic mobility shift assays (EMSAs) (43). Using these data to guide us, along with the fact that CAP is a DNA-binding transcription factor, we hypothesized that CAP directly bound to P*_lux_* to regulate transcription. Although the Chatterjee study showed that CAP could bind P*_lux_* DNA (∼400-500-bp fragment), a precise binding site was never identified and controls to determine DNA-sequence specificity were not performed (43). Thus, we aimed to determine whether CAP specifically interacted with the *lux* promoter DNA. We began with a bioinformatics approach to predict *V. campbellii* CAP binding sites within P*_lux_* using published consensus sequences from *V. cholerae* CAP (94% amino acid identity) and *E. coli* CAP (99% amino acid identity) (38, 66). Using the *E. coli* consensus, MEME Suite-FIMO (67) identified two possible CAP binding sites within P*_lux_* when restricted to a *p*-value of 0.05. These binding sites are centered at approximately −329 and −246 from the *luxCDABE* transcription start site (TSS) and named CAP binding sites 1 and 2, respectively (Fig. 4A, Table S1). Using the *V. cholerae* consensus and a *p*-value restriction of 0.05, MEME Suite-FIMO (67) again identified CAP binding site 2 as well as a binding site centered approximately −67 from the *luxCDABE* TSS, named CAP binding site 3 (Fig. 4A, Table S2). We focused on predicted CAP sites 1, 2, and 3 for further analysis.

**Figure 4.**
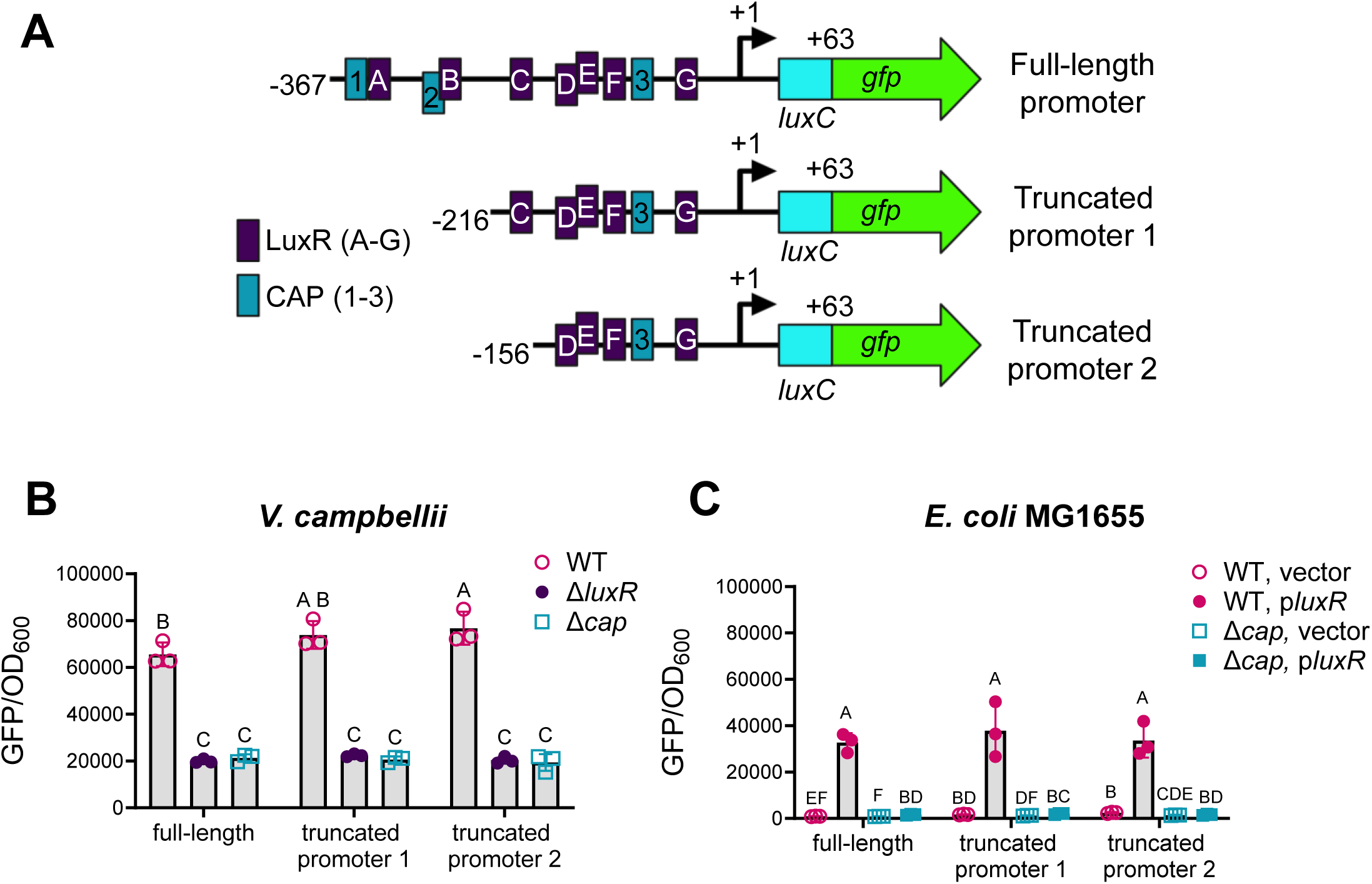
CAP and LuxR are required for *lux* transcription in *V. campbellii* and *E. coli*. (A) Diagram of the full-length *luxCDABE* promoter region (−367 to +63), truncated promoter 1 (−216 to +63), and truncated promoter 2 (−156 to +63) transcriptional *gfp* fusions. LuxR binding sites (labeled A-G) and predicted CAP binding sites (labeled 1-3) are represented as dark purple and teal boxes, respectively. (B) Fluorescence production (GFP/OD_600_) was measured in wild-type (WT), Δ*luxR*, and Δ*cap V. campbellii* strains harboring either a plasmid with the full-length promoter (pCS019), truncated promoter 1 (pAB101), or truncated promoter 2 (pAB102) *gfp* fusions. A two-way ANOVA was performed on normally distributed data (Shapiro-Wilk test) followed by Tukey’s multiple comparisons test (*p*<0.05; *n*=3). (C) Fluorescence production (GFP/OD_600_) was measured in WT and Δ*cap E. coli* MG1655 strains harboring either 1) an empty vector control (pLAFR2) or a plasmid expressing *luxR* from its native promoter (pKM699) and 2) a plasmid with either the full-length promoter (pCS042), truncated promoter 1 (pAB118), or truncated promoter 2 (pAB119) *gfp* fusions. A two-way ANOVA test was performed on normally distributed log transformed data (D’Agostino-Pearson test on the residuals) followed by Tukey’s multiple comparisons test (*p<*0.05; *n*=3). (B,C) Different letters above bars indicate differences in significance between strains in pairwise comparisons. Error bars represent the standard deviation.

### Analysis of predicted CAP binding sites on P*_lux_* transcription

To test if predicted CAP binding sites were necessary for P*_lux_* transcription, we constructed promoter truncations. We reasoned that if we disrupted an important activating site, we would see a reduction in transcription. Because bioluminescence is sensitive to many factors like oxygen levels, pH, and accumulation of the proper fatty aldehyde substrate, a change in light levels could be due to an effect of the bioluminescence mechanism rather than a reduction in transcription (58, 60). Thus, we used P*_lux_* transcriptional fusions to *gfp* to assay the impact of CAP and LuxR on P*_lux_* transcription (Fig.4A). When testing the full-length promoter construct (−367 to +63), we observed high fluorescence in a WT strain that decreased in both Δ*luxR* and Δ*cap* strains, indicating that CAP and LuxR are both required for P*_lux_* transcription (Fig. 4B). Next, we constructed promoter truncations of various lengths to test if the bioinformatically predicted CAP binding sites were necessary for transcription (Fig. 4A). These assays also tested the importance of previously validated LuxR binding sites A-G on P*_lux_* transcription (Fig. 4A) (47, 49). Truncated promoter 1 (−216 to +63) and truncated promoter 2 (−156 to +63) were both sufficient to drive GFP production, indicating that CAP binding sites 1 and 2 and LuxR binding sites A-C were not necessary for transcription at P*_lux_* (Fig. 4B). Previous reports from our group have demonstrated that LuxR sites D, E, F, and G are required for transcriptional activation, thus we did not construct truncations past site D (45).

Next, we tested our P*_lux_*-*gfp* constructs in *E. coli* MG1655 to determine if CAP and LuxR are both required for transcriptional activation in a heterologous system. Here, we transformed *E. coli* with two plasmids: the first containing *V. campbellii luxR* under expression from its own promoter and the second containing P*_lux_* truncations fused to *gfp*. Fluorescence was observed only when LuxR and CAP were both present (Fig. 4C), indicating that even in a heterologous system, both CAP and LuxR are necessary for P*_lux_* transcription. Importantly, these results also suggest that *E. coli* CAP can cross-complement *V. campbellii* CAP and that CAP acts directly at the *lux* promoter. From these collective data, we predicted that CAP bound to binding site 3 to regulate P*_lux_* transcription in *V. campbellii* and *E. coli*.

### CAP does not bind P_lux_ *in vivo*

From the results of our P*_lux_* truncation experiment, we hypothesized that CAP binds P*_lux_ in vivo* to regulate *lux* transcription. To test our hypothesis, we used Chromatin Immunoprecipitation coupled with high-throughput Illumina sequencing (ChIP-seq) to identify CAP binding sites *in vivo.* We used commercially available monoclonal *E. coli* CAP antibodies (99% amino acid identity) that we showed are effective via western blot (Fig. 2F) and that were used for CAP ChIP-seq in *V. cholerae* (38, 68). We assayed CAP binding at high cell density (OD_600_ = 1.0) when light production is maximal, and we observed 479 peaks that were at least 2-fold enriched compared to the input control (*q*-value < 0.01) in a WT background (Fig. S3A, Dataset S3). We examined CAP binding at predicted (adenylate cyclase; *cyaA*) and non-predicted (RNA polymerase subunit D; *rpoD*) promoter regions, which were predicted using published CAP ChIP-seq datasets from *E. coli* and *V. cholerae* (38, 69). We found that CAP bound at the *cyaA* promoter (Fig. 5B) and did not bind at the *rpoD* promoter (Fig. 5C). We further validated these results using ChIP-qPCR and found that the *cyaA* promoter was highly enriched in our pulldown and the *rpoD* promoter was not (Fig. 5D). From our two validation methods, we concluded that our ChIP-seq was a robust and accurate method to identify CAP binding locations across the *V. campbellii* chromosome. However, under the conditions we tested, we were unable to identify an *in vivo* CAP binding site within P*_lux_* (Fig. 5A).

**Figure 5.**
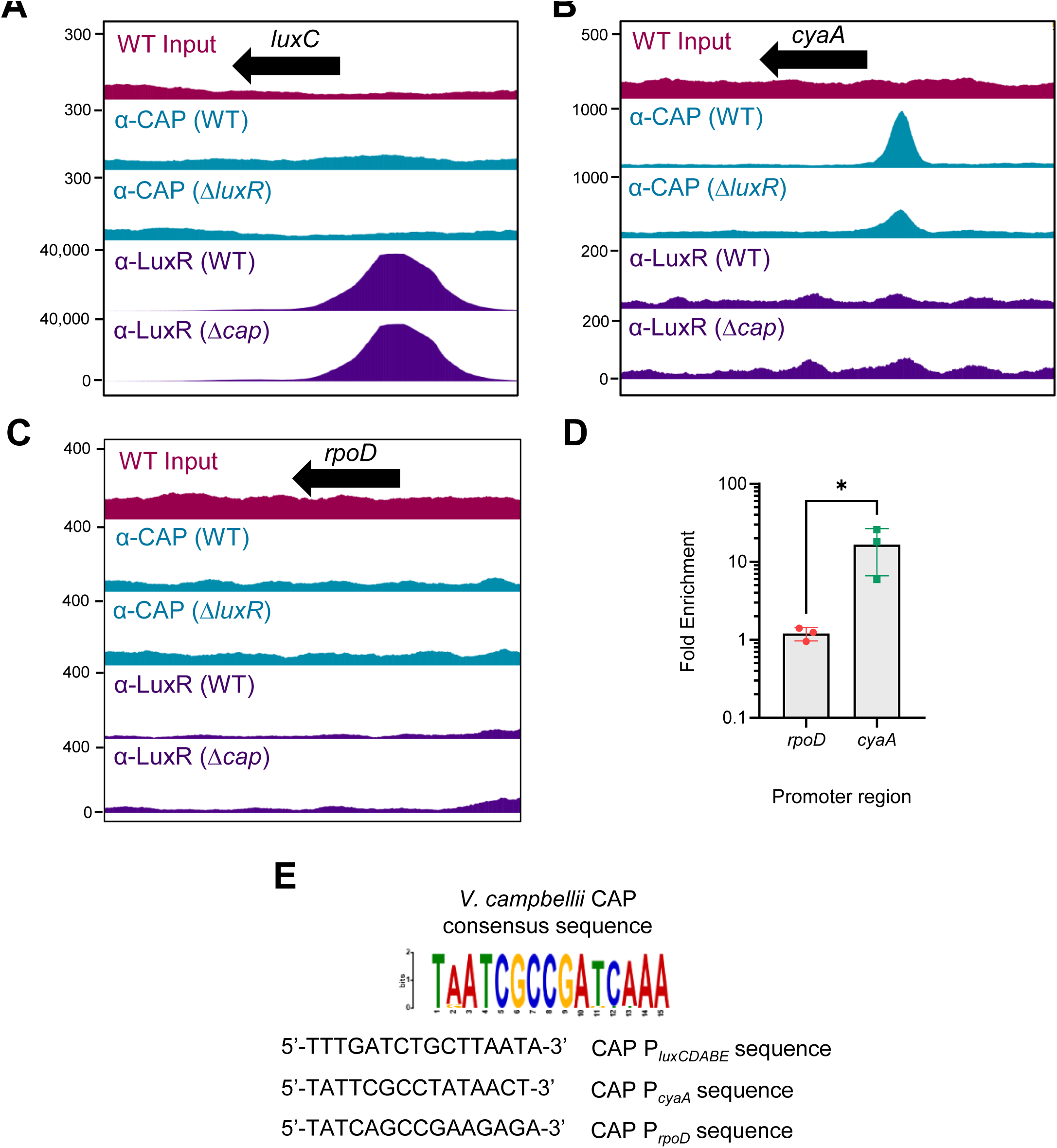
Analysis of CAP and LuxR binding peaks. (A-C) ChIP-seq was performed in WT, Δ*cap*, and Δl*uxR V. campbellii* strains with either α-CAP or α-LuxR antibodies at an OD_600_ of 1.0. The top pink panel is the sequenced WT input DNA that was untreated with antibodies. The ChIP-seq binding profiles of CAP in a WT strain and a Δ*luxR* strain are shown in teal as the second and third panels from the top, respectively. The binding profiles of LuxR in a WT strain and a Δ*cap* strain are shown in purple as the fourth and fifth panels from the top, respectively. The *luxC* gene (A), the *cyaA* gene (B), and the *rpoD* gene (C) are indicated as black arrows. Peak height (DNA enrichment) is displayed as the y-axis. A representative image from two biological replicates was chosen for the figure. Screenshots of ChIP peaks were captured using JBrowse 2. (D) ChIP was performed in WT *V. campbellii* strains using α-CAP antibodies. Primers were designed to amplify the promoters of a negative (*rpoD*) and a positive (*cyaA*) target from the extracted ChIP DNA using qPCR. Fold enrichment represents IP DNA normalized to input DNA for each target and was measured using the ΔΔC_T_ method and *hfq* as an internal control. The asterisk above the bar indicates significance, as determined using a lognormal Welch’s *t*-test (*p<*0.05; *n*=3) performed on lognormally distributed data (Shapiro-Wilk test). Error bars represent the standard deviation. (E) All significant CAP ChIP peaks were extracted and then run through MEME-ChIP to identify a consensus sequence (shown as a colorful logo map). The CAP consensus sequence was then run through MEME-Suite FIMO to identify potential CAP binding sites within P*_luxCDABE_*, P*_cyaA_* (positive control), and P*_rpoD_* (negative control). The sequences that were identified using MEME-Suite FIMO are listed 5’-3’ and additional information can be found in supplemental table 3.

One hypothesis that could explain these results was that LuxR and CAP may compete for binding at the *lux* locus because there are seven LuxR binding sites within P*_lux_* (Fig. 4A) and LuxR protein levels are higher than CAP levels at an OD_600_ of 1.0 in a WT strain (Figs. 2E, 2F, S6). To test this hypothesis, we performed ChIP-seq under the same conditions in a Δ*luxR* strain to see if CAP can bind P*_lux_* when LuxR is absent. We identified 198 CAP peaks that were at least 2-fold enriched compared to the input control (*q*-value < 0.01) in the Δ*luxR* background. We note that there were fewer CAP peaks and overall enrichment observed in the Δ*luxR* background compared to the WT background, indicating that LuxR may influence CAP binding at many sites across the chromosome (Fig. S3A, Dataset S3). However, we were still unable to identify a CAP binding site within P*_lux_* (Fig. 5A).

**Figure 6.**
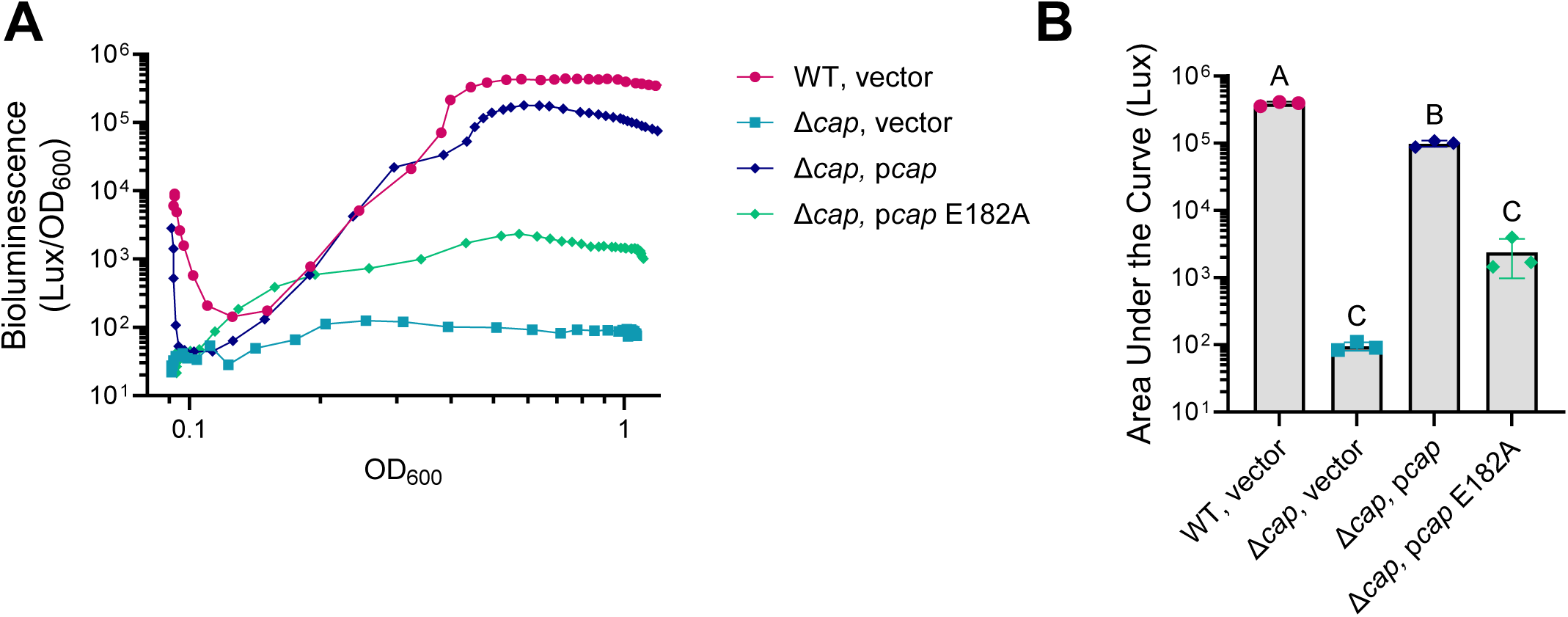
CAP DNA binding ability is necessary for bioluminescence production. (A) Bioluminescence production and OD_600_ were measured over 24 hours for Wild-type (WT) and Δ*cap V. campbellii* strains harboring either an empty vector (pCS027) or an episomal plasmid expressing *cap* (pAB100) or *cap* E182A (pCM026) from an IPTG-inducible promoter (25 μM). The figure is a representative graph from three biological replicates and each line on the graph is an average of two technical replicates. (C) Area under the curve (Lux) was measured from bioluminescence vs OD_600_ plots for three biological replicates using the same strains and conditions from (B). A one-way ANOVA test was performed on normally distributed data (Shapiro-Wilk Test), followed by Tukey’s multiple comparisons test between strains. Different letters above bars indicate differences in significance between strains in pairwise comparisons (*p*<0.05; *n*=3). Error bars represent the standard deviation. Additional replicates with IPTG titrations can be found in supplemental figure 4.

Because CAP binding site 3 was predicted using the *V. cholerae* CAP consensus sequence, we re-ran our analysis searching for predicted CAP binding sites within P*_lux_* using our MEME-ChIP (70) generated *V. campbellii* CAP consensus sequence (Fig. 5E, Table S3). Using MEME Suite-FIMO (67), we re-identified potential CAP binding site 3 with a p-value of 0.00261 (Fig. 5E, Table S3). To gauge the significance of CAP site 3, we then ran MEME Suite-FIMO (67) to identify CAP binding sites within known CAP regulated promoters like *cyaA* (Figs. 5B, 5D, 5E, Table S3, *p*-value: 0.000621) and promoters where CAP does not bind like within *rpoD*, (Figs. 5C, 5D, 5E, Table S3, *p*-value: 0.00261) to set parameters for our analysis. Notably, the *p*-value for CAP binding site 3 was identical between P*_lux_* and our negative control *rpoD,* suggesting that predicted CAP site 3 was a false positive result (Table S3). Taken together with our *in vivo* ChIP-seq data, we concluded that CAP does not bind P*_lux_* in *V. campbellii*.

### CAP does not influence LuxR *in vivo* DNA binding activity at P*_lux_*

Because we have shown that CAP does not bind P*_lux_* and does not affect LuxR levels, but still controls *luxCDABE* transcription, we hypothesized that perhaps CAP affects LuxR DNA binding activity. Thus, the lack of bioluminescence in a Δ*cap* strain could be explained by LuxR failing to bind P*_lux_* to activate transcription. To test this hypothesis, we performed ChIP-seq at high cell density (OD_600_ = 1.0) when light production is maximal and used polyclonal LuxR antibodies (71) to pull down LuxR in WT, Δ*cap*, and Δ*luxR* strains. We validated our native LuxR ChIP-seq results by comparing them to our previously published ChIP-seq using FLAG-LuxR (47, 49) and observed similar binding profiles across the chromosome (Dataset S3). We observed 476 peaks in a WT background and 640 peaks in a Δ*cap* background that were at least 2-fold enriched compared to the input controls (*q*-value < 0.01) (Dataset S3, Fig. S3B). We identified >200 new LuxR peaks in the Δ*cap* background, suggesting that CAP influences where LuxR can bind across the chromosome. However, we note that overall enrichment across the chromosome looks similar between the two backgrounds (Fig. S3B). We also note that there is some noise in the Δ*luxR* background control (Fig. S3B), which could be due to the slightly nonspecific binding nature of the LuxR polyclonal antibodies. When we checked P*_lux_* for the presence of LuxR binding in the Δ*cap* strain, we observed that LuxR was indeed binding and there were no detectable differences in peak height between WT and Δ*cap* backgrounds (Fig. 5A). From these data, we concluded that LuxR DNA binding activity is not affected by CAP at P*_lux_*.

### CAP DNA binding activity is necessary for bioluminescence

Because CAP was not binding to P*_lux_* or affecting LuxR DNA binding activity, we hypothesized that either 1) CAP was regulating another protein (an intermediate factor) that is necessary for bioluminescence or 2) a CAP-LuxR interaction was necessary for transcriptional activation. Indeed, CAP and LuxR homologs interact in *V. cholerae* and data suggest they interact in *V. vulnificus* (37, 40, 41). Additionally, we pulled down CAP with LuxR in a previous co-immunoprecipitation experiment (50). The CAP E181 residue in *E. coli* (E182 in *V. campbellii*) is located within the DNA-binding domain and was demonstrated to be important for *E. coli* CAP DNA recognition (18). Thus, to test the role of CAP DNA binding, we mutated *V. campbellii* CAP at residue E182 to an alanine (E182A). We placed the *cap* mutant under an IPTG-inducible promoter (P*_tac_*) and measured bioluminescence in a Δ*cap* strain. We determined that a CAP E182A mutant was deficient in bioluminescence production like a Δ*cap* strain (Fig. 6A, 6B, S4A, S4B). However, when we measured *luxC* transcript levels at an OD_600_ of 1.0, we observed that transcription was reduced compared to WT but not fully ablated, like the Δ*cap* strain (Fig. S4C). We observed similar results when measuring transcript levels for a known CAP-regulated gene *cyaA* (Fig. S4D), indicating that additional residues beyond E182 may be necessary for DNA recognition. Regardless, our data support a role for CAP DNA recognition and the CAP E182 residue in regulation of bioluminescence production, likely through regulation of an intermediate factor.

### LuxR and CAP directly co-regulate 11 promoters

Although the *luxCDABE* genes were the most highly co-regulated genes by CAP and LuxR, we sought to determine if other co-regulated genes were directly regulated via binding at the promoter by one or both proteins. We first searched for genes directly regulated by CAP and LuxR by comparing our ChIP-seq and RNA-seq datasets. The parameters for our analysis were 1) genes must be strongly regulated by CAP or LuxR (≥4-fold regulated) and 2) genes must possess a ≥2-fold enriched CAP or LuxR ChIP peak within the promoter. Using these criteria, we found that CAP directly regulated 206 promoters and LuxR directly regulated 36 promoters (Fig. 7F, Dataset S4). We next sought to determine if CAP and LuxR were both at the promoters for genes we identified as being co-regulated by CAP and LuxR (Figs. 1E, 1F). Since our direct regulon analysis had stringent parameters that overlooked lowly expressed genes identified in our original RNA-seq analysis, we manually inspected the regulons to ensure we captured all directly regulated loci in our analysis. We found 5 additional promoters regulated by LuxR that were initially overlooked, bringing the total number of promoters regulated by LuxR to 43. We found that CAP and LuxR both bound to 11 co-regulated promoters, LuxR bound without CAP to 32 co-regulated promoters, and CAP bound without LuxR to 7 co-regulated promoters (Fig. 7F, Dataset S4).

**Figure 7.**
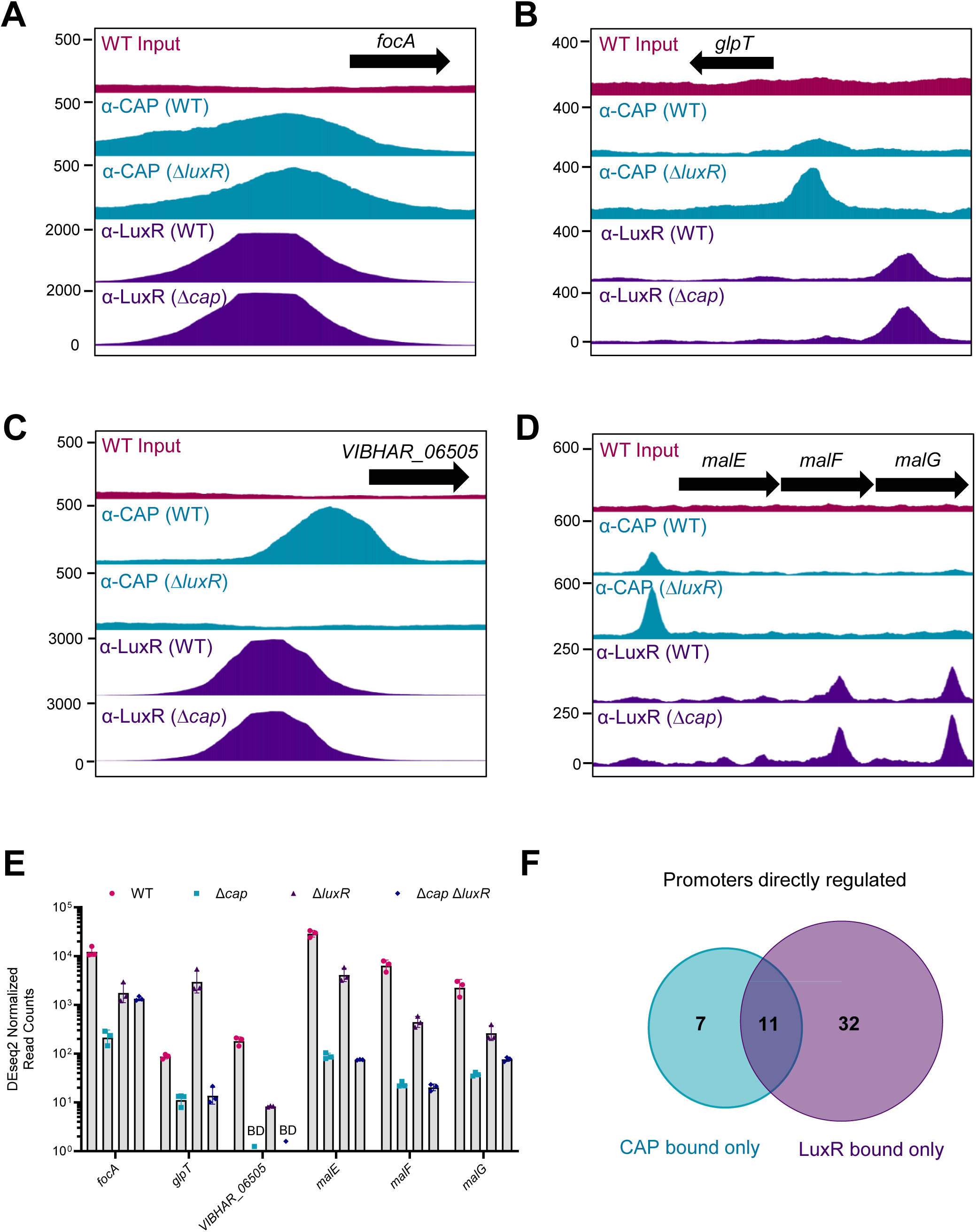
CAP and LuxR use multiple mechanisms to co-regulate genes. (A-D) CAP and LuxR binding profiles for directly co-regulated genes *focA* (A), *glpT* (B), *VIBHAR_06505* (C), and *malEFG* (D) at an OD_600_ of 1.0. The ChIP-seq binding profiles of CAP in a WT strain and a Δ*luxR* strain are shown in teal as the second and third panels from the top, respectively. The binding profiles of LuxR in a WT strain and a Δ*cap* strain are shown in purple as the fourth and fifth panels from the top, respectively. The top pink panel is the sequenced WT input DNA that was untreated with antibodies. Genes are indicated as black arrows. Peak height (DNA enrichment) is displayed as the y-axis. Screenshots of ChIP peaks were captured using JBrowse 2 version 3.6.4. (E) The normalized transcript counts for the genes indicated in A-D as determined from an RNA-seq analysis that was performed on RNA collected from WT, Δ*cap,* Δ*luxR,* and Δ*cap* Δ*luxR V. campbellii strains* grown to high cell density (OD_600_ of 1.0). Error bars represent standard deviation. BD=below detection. (F) ChIP-seq and RNA-seq datasets were compared to find genes that were directly regulated using the following criteria: genes were 1) ≥ 4-fold regulated by CAP and LuxR and 2) contained a ≥ 2-fold enriched CAP and/or LuxR binding site within the promoter. The instances of CAP binding promoters only is represented in teal, and the instances of LuxR binding promoters only is represented in purple. The instances of both CAP and LuxR binding promoters is represented by the overlap between the two circles.

At the directly co-regulated promoters, we discovered several distinct patterns of binding and transcriptional effects. At the *focA* (formate transporter) promoter, CAP and LuxR were both present regardless of the presence or absence of the other protein and their binding peaks directly overlapped (Fig. 7A). The DEseq2 normalized read counts from our RNA-seq analysis (referred to hereafter as normalized reads) indicated that CAP and LuxR both activated *focA* expression with different strengths, suggesting a synergistic co-regulation strategy (Fig. 7E, Dataset S1). A variation of this pattern also occurred at the *glpT* (glycerol-3-phosphate transporter) promoter, where CAP and LuxR were both present regardless of the presence or absence of the other protein (Fig. 7B). However, both proteins displayed taller binding peaks in the absence of the other, suggesting that there may be a competition for binding. The normalized reads indicated that CAP activated *glpT* expression while LuxR repressed expression, suggesting an antagonistic co-regulation strategy (Fig 7E, Dataset S1). At the *VIBHAR_065505* promoter (putative virulence operon), CAP only bound if LuxR was also present, but LuxR bound in the presence or absence of CAP, suggesting that an interaction between CAP and LuxR may be necessary for CAP binding (Fig. 7C). The normalized reads showed that CAP and LuxR were both activators of the *VIBHAR_065505* promoter (Fig. 7E, Dataset S1). We also observed instances of more complicated regulation, like at the *malEFG* (maltose utilization) locus, where CAP bound the promoter and LuxR bound within *mal* genes possibly to either terminate transcription or regulate an intragenic small RNA or small protein (Fig. 7D). The *malEFG* locus was counted as bound by CAP only since our criteria only included binding sites within promoters (Fig. 75FF). The normalized reads showed that CAP was the stronger activator of the *malEFG* locus while LuxR had a smaller effect on transcriptional activation (Fig. 7E, Dataset S1). From these data, we predicted that CAP and LuxR directly co-regulate transcription at a dozen promoters using several putative mechanisms, including synergistic binding, competitive binding, and physical interactions.

## Discussion

CAP and LuxR homologs have been studied extensively for decades for their responses to nutrient starvation and population levels, respectively. However, how these transcription factors intersect across the chromosome to globally co-regulate gene expression in *Vibrio campbellii* was unknown. In this study, we used ChIP-seq and RNA-seq to define individual and combined regulons for CAP and LuxR. We found that CAP and LuxR utilize several strategies-both direct and indirect- to co-regulate over 100 genes in *V. campbellii.* At the promoters for co-regulated genes, we either found both CAP and LuxR bound, only CAP or only LuxR bound, or neither protein bound (Fig. 7F). When both proteins were bound, we observed that CAP and LuxR co-activated gene expression by synergistically binding promoters, antagonized gene expression through a competitive binding mechanism, and potentially interacted with each other for maximal gene expression (Fig. 7). We note that CAP and LuxR each control dozens of transcription factors in their individual and combined regulons (Dataset 1). Thus, we speculate that a major mechanism of co-regulation is through the regulation of additional transcription factors that directly co-regulate genes with CAP or LuxR. We hypothesize this is the mechanism of regulation occurring at co-regulated promoters that have either CAP only, LuxR only, or neither protein bound (Fig. 7F, Dataset S4). Intriguingly, CAP and LuxR appear to influence global binding of the other protein. We found that CAP bound to fewer loci in the absence of LuxR suggesting LuxR has a positive influence on CAP binding at hundreds of loci, possibly through a protein-protein interaction (Dataset S3). Additionally, CAP appears to antagonize or modulate LuxR binding because in the absence of CAP, LuxR bound >200 additional loci (Dataset S3). This study validated several existing connections between cAMP sensing and quorum sensing (QS) as well as established many new connections. The finding that cAMP signaling and QS signaling intersect at numerous loci in *V. campbellii* and *V. cholerae* (37) lends support to the hypothesis that these two systems are widely used by vibrios in nature to efficiently respond to their diverse and fluctuating environments.

CAP has been previously implicated in the regulation of specific QS genes, like regulation of the CAI-1 autoinducer synthase (*cqsA*) in *V. cholerae* (30, 35). In alignment with these findings, we found that CAP strongly activated expression of *cqsA* in *V. campbellii* (Fig. S1, Dataset S1). In *V. cholerae*, it was proposed that CAP controls *cqsA* expression through a post-transcriptional mechanism (35), but because we measured *cqsA* transcript levels using RT-qPCR (Fig. S1) in our study, we cannot yet report how CAP controls *cqsA* expression in *V. campbellii*. However, we found that *cqsA* was strongly co-regulated by both CAP and LuxR, but only LuxR bound the promoter, which could support a role for CAP in post-transcriptional regulation (Datasets S1, S3, S4). CAI-1 is sensed at the membrane by its cognate receptor CqsS and the *cqsA* and *cqsS* genes are next to each other on the chromosome but are oriented opposite directions and each have their own promoter (72–75). Upon closer examination of our ChIP-seq data, we identified that CAP and LuxR are both present at the *cqsS* promoter, but they do not affect *cqsS* transcription, thus this promoter was overlooked in our direct regulon screen because it did not meet our criteria (Datasets S1-S3). Perhaps CAP and LuxR binding at the CqsS promoter influences *cqsA* expression rather than *cqsS* expression? Indeed, CAP has a complex relationship with autoinducers in *V. campbellii*. One of the strongest CAP peaks we consistently identified was within the gene for *luxM*, the HAI-1 autoinducer synthase (Dataset S3). However, we did not identify *luxM* as being a CAP regulated gene (Dataset S1). Could the peak within *luxM* be important for terminating transcription or perhaps regulating an intragenic small RNA (sRNA) or small protein? This phenomenon of CAP binding within genes has also been documented in *E. coli* (69). Additionally, CAP is known to regulate several sRNAs in different species and in *E. coli*, CAP regulates a sRNA that regulates production of AI-2 (76–81). Furthermore, recent research has shown that CAP and LuxR co-regulate an intragenic sRNA important for phage production in *V. cholerae* (39). The position of the CAP peak within *luxM* is intriguing but more work would need to be done to validate if this binding site is relevant and impacts HAI-1 production. In this study, we measured LuxR levels—which indirectly reported on AI levels since they are directly correlated—and we found no difference between LuxR levels in Δ*cap* and WT strains (Fig. 2E), indicating that CAP does not influence AI levels under the conditions we tested. AIs are thought to turn on the QS system at different times during the growth curve, and it was previously proposed that CAI-1 operates at very low cell densities in *V. campbellii* but at high cell densities in *V. cholerae* (72, 73, 82). Thus, it is possible that CAP does influence CAI-1 levels at different stages during growth, but this was not measured in our assays. We speculate that CAP regulation of *cqsA* could be important under different conditions, such as during colonization or in low cell density states.

For decades, bioluminescence has been reported as a density-dependent QS regulated phenotype. While this is true, we have shown that both QS (via LuxR) and nutrient sensing (via cAMP-CAP) are required for *lux* expression, which has also been demonstrated in other bioluminescent organisms like *A. fischeri* (62–65). Interestingly, we found that we can modulate the timing of *lux* expression (and bypass canonical density-dependence) through growth in glucose or by controlling either the amount of CAP, cAMP, and/or LuxR (Figs. 2A, 3A). Most of the field’s knowledge about the connection between CAP and carbon sources comes from *E. coli*, where cyclic-AMP is produced in response to low preferred carbon sources like glucose (23, 83, 84). Thus, when glucose is high, intracellular cAMP is low, and we reasoned that cAMP-regulated phenotypes (like bioluminescence) should also be low/off in glucose. We were surprised to find that bioluminescence was brightest and began the earliest when grown in minimal media with glucose (Figs. 3A, 3B). This finding was contrary to studies in *A. fischeri* where glucose repressed bioluminescence production through carbon catabolite repression via CAP (62–65). This suggests that perhaps the glucose utilization pathway is different in *V. campbellii* compared to the canonical model in *E. coli.* Intriguingly, in *V. cholerae*, glucose represses CAI-1 production, which then represses HapR production and HapR regulated-phenotypes (30, 35). These findings are contradictory to what we observed here in *V. campbellii* BB120, and future work should be done to test LuxR and CAI-1 levels in media with glucose to examine the impact of glucose on QS.

It has been documented for decades that *Vibrio* species can respond to extracellular cAMP through an unknown mechanism, and we confirmed these findings here in *V. campbellii*. We note that *V. campbellii* is sensitive to the concentration of extracellular cAMP and can modulate cAMP-regulated phenotypes accordingly. For instance, the timing and output of bioluminescence was different in media with no cAMP, residual amounts of cAMP, and 1 mM extracellular cAMP (Figs. 3A, 3B). Because many living organisms use cAMP as a signaling molecule, we propose that in addition to being an intracellular signaling molecule communicating the nutritional state of the cell, cAMP could also serve as a host-sensing molecule, allowing vibrios to fine-tune their transcription depending on their environment. We cannot overlook the fact that *V. cholerae* produces the cholera toxin, which overstimulates host cAMP production (85). Thus, we postulate that the ability to sense different amounts of extracellular cAMP is beneficial for *V. campbellii*. Beyond CAP, cAMP, and glucose, there are several instances of nutrient regulators intersecting with QS in *Vibrio campbellii*. For instance, our group recently showed that QS via LuxR regulates methionine and tetrahydrofolate biosynthesis genes (86). Additionally, previous work has shown that homocysteine and MetR co-repress bioluminescence through regulation of LuxR levels (43). Lastly, L-arginine has been reported to stimulate bioluminescence for decades through an unknown mechanism (87–89). Clearly, *Vibrio* bacteria use multiple nutrient signals to regulate bioluminescence and LuxR levels, likely for proper timing of bioluminescence genes since they are expensive for cells to produce.

Currently, our data support a model where CAP controls an intermediate factor that is necessary for transcription of bioluminescence genes. While we did not search for the intermediate factor in this study, we note that the factor must also exist in *E. coli* since deletion of *cap* resulted in a loss of transcriptional activation at the *lux* promoter in the heterologous system (Fig. 4C). We hypothesize that CAP directly regulates transcription of the intermediate factor because a loss in CAP DNA binding activity resulted in a loss of bioluminescence (Fig. 6). To fit our model, CAP could either activate an activator or repress a repressor. We speculate that the identity of the intermediate factor could be a transcription factor required for activation or repression of *lux* transcription, a small regulatory RNA (sRNA) or RNA binding protein that either stabilizes or destabilizes *lux* mRNA, or a post translational modifier of LuxR that does not affect DNA binding ability but affects LuxR’s ability to either stimulate transcription or make P*_lux_* more accessible for transcriptional activation by removing anti-activators such as H-NS (49). We found 206 promoters that were directly regulated by CAP and identified numerous candidates for our intermediate P*_lux_* regulator, including several predicted transcription factors, acetyltransferases, and RNA binding proteins (Dataset S1, S4). While it is possible that the P*_lux_* regulator is one of the hundreds of genes we identified, there are simply too many genes to screen each one individually. A more targeted approach, like a screen or a selection, would be more efficient at finding the intermediate regulator. The screen should be designed in a way that allows for finding factors that affect transcription rather than bioluminescence production.

We propose an updated model of CAP and LuxR co-regulation of the bioluminescence operon in Fig. 8. In a Δ*cap V. campbellii* strain, LuxR is produced and binds across the *lux* promoter. However, *lux* genes are not expressed and bioluminescence is off (Fig. 8A). In a WT strain, LuxR binds across the promoter and CAP controls production of an intermediate factor that could either regulate LuxR activity (but not DNA binding) via a post-translational modification or ligand, directly regulate transcription by binding the *lux* promoter, or regulate translation of *lux* mRNA (Fig. 8B). Ultimately, this study advances our knowledge about CAP and LuxR global co-regulation in *Vibrio campbellii.* Our findings here help contextualize how *Vibrio* bacteria sense and respond to their environments using populational signals and nutrient availability cues. Our data add support to the growing hypothesis in the field that CAP and LuxR-homolog global co-regulation is a conserved mechanism used by *Vibrio* species to sense and respond to their environments.

**Figure 8.**
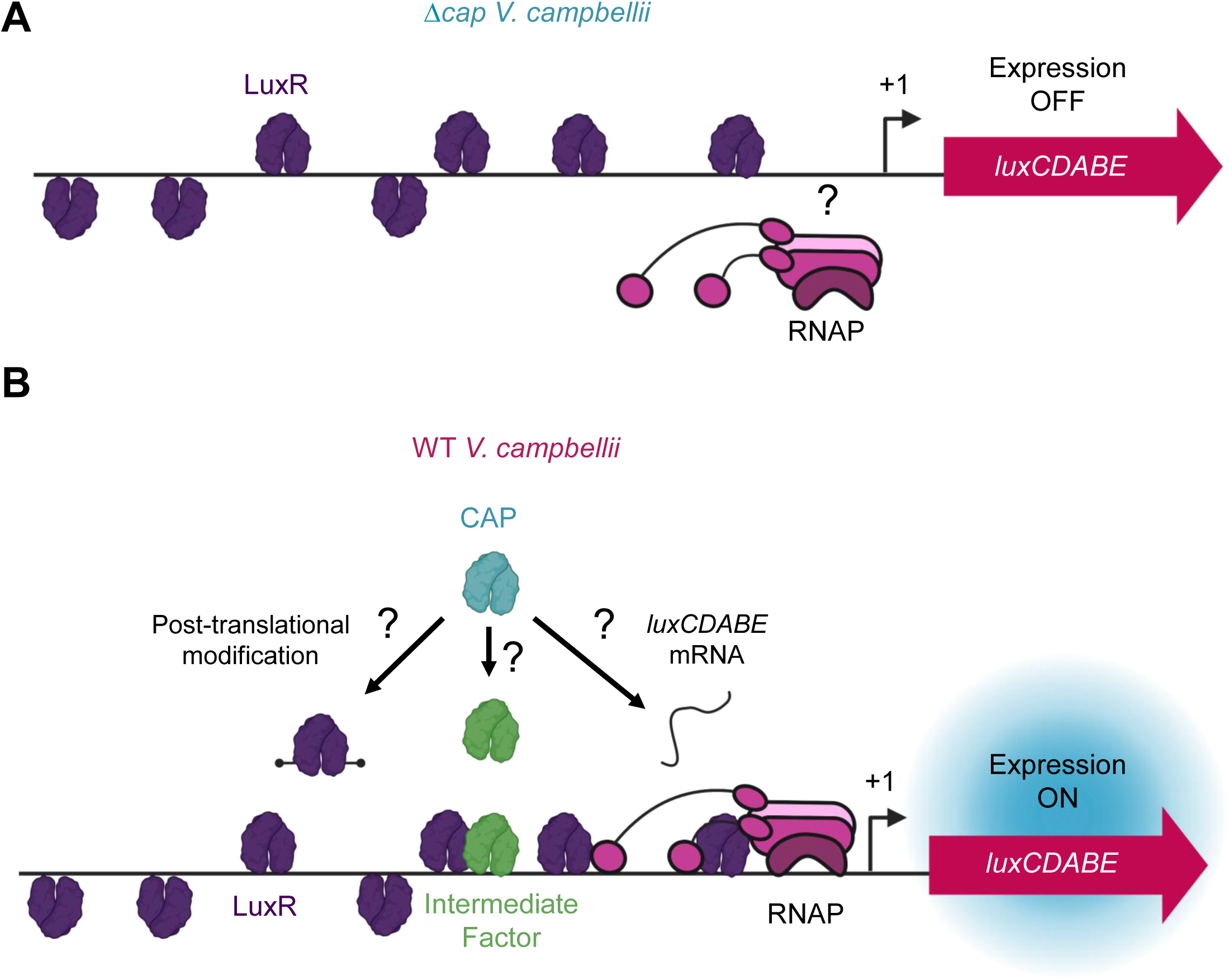
Model of CAP and LuxR co-regulation of P*_lux_*. (A) In a Δ*cap V. campbellii* strain, LuxR binds across the promoter, but *luxCDABE* expression is off and bioluminescence is not produced. (B) In a WT *V. campbellii* strain, CAP and LuxR strongly co-regulate *luxCDABE* expression but only LuxR binds the promoter. CAP controls an intermediate factor that either post-translationally modifies LuxR, binds the *luxCDABE* promoter itself, or influences the stability of *luxCDABE* mRNA. RNAP= RNA Polymerase. Question marks indicate open hypotheses that we did not rule out in this study. The model was created in BioRender (https://BioRender.com/op4mmsg).

## Materials and Methods

### Bacterial Strains and Media

*Vibrio campbellii* BB120 (a.k.a. ATCC BAA-1116; previously classified as *V. harveyi* BB120) (13) and derivatives were grown at 30°C shaking (250-275 RPM) in lysogeny broth marine medium (LM; lysogeny broth (LB) supplemented with an additional 10 g NaCl/L) or M9 Minimal Medium supplemented with 20 mM glucose. When selection was required, *V. campbellii* culture medium was supplemented with either 100 µg/mL kanamycin, 10 µg/mL chloramphenicol, 100 µg/mL gentamycin, and/or 50 U/mL polymyxin B. Plasmids were electroporated into and stored in either *Escherichia coli* S17-1λpir or *E. coli* BL21(DE3) cells and their derivatives. All *E. coli* strains were grown in LB at 37°C shaking (250-275 RPM); when selection was required, *E. coli* culture medium was supplemented with 40 µg/mL kanamycin, 10 µg/mL chloramphenicol, 100 µg/mL gentamycin, 10 µg/mL tetracycline, and/or 100 µg/mL ampicillin. Plasmids were transferred to *V. campbellii* via conjugation and exconjugants were selected using polymyxin B. Where applicable, IPTG was added to a final concentration between 1 μM-25 μM to induce gene expression, and the specific concentration is noted in figure captions. Cyclic-AMP (cAMP) was dissolved in 50 mM Tris pH 8.0 and added to a final concentration of 1 mM for the applicable assays. All strains used in this study are listed in Table S4.

### Molecular Methods

PCR and site-directed mutagenesis were performed using Phusion HF polymerase (New England Biolabs; NEB) or iProof HF DNA Polymerase (BioRad). All restriction enzymes were purchased from NEB and used according to the manufacturer’s protocols. All oligonucleotides used for cloning, site-directed mutagenesis, qPCR, and RT-qPCR were purchased from Integrated DNA Technologies. Cloning primers are available upon request while RT-qPCR and qPCR primers are listed in Table S6. DNA samples were resolved using 0.8%-1% agarose gels dissolved in 1xTBE. All genome and plasmid genotypes were confirmed via sanger and whole-plasmid sequencing, respectively, at Eurofins Genomics. All plasmids used in this study are listed in Table S5. All gene names and locus tags used in this study are located in Table S7.

### Cloning procedures are available upon request

#### Deletion and Knockdown Strain Construction

All deletion derivatives of *V. campbellii* BB120 were constructed as previously described (45) using the suicide plasmid (pRE112) and confirmed via sequencing. To construct complement strains, we cloned the gene of interest under an IPTG-inducible promoter onto a plasmid containing Tn7 integration sites (pJMP1339) that insert the plasmid onto the chromosome downstream of the conserved *glmS* gene and confirmed via sequencing (90–92).

#### RNA extraction, RNA-seq, and Reverse Transcriptase quantitative PCR (RT-qPCR)

Overnight cultures were back-diluted and grown in fresh LM media to an OD_600_ of 1.0 and 3.0 ODs were collected to extract RNA for further analysis. RNA was extracted using a Trizol/chloroform method and RT-qPCR was performed using a SensiFAST SYBR Hi-ROX One-Step Kit as previously described by Simpson and colleagues (93). Primers for RT-qPCR are in Table S6. All RT-qPCR experiments were performed with 3 biological replicates and 2 technical replicates using *hfq* as an internal control and the ΔΔ*C_T_* method for analysis. Extracted RNA was submitted to the Indiana University Center for Genomics and Bioinformatics (CGB) for library preparation (Illumina TruSeq Stranded mRNA), ribosomal RNA depletion (Illumina Ribo-Zero Plus rRNA depletion), and sequencing (NextSeq Illumina 1000/2000).

#### RNA-seq Analysis

Reads were adapter trimmed and quality filtered using Trimmomatic ver. 0.38 (94) setting the cutoff threshold for average base quality score at 20 over a window of 3 bases. Reads shorter than 20 bases post-trimming were excluded. About 93% of the reads have both the mates passing the quality filters. Reads were mapped to the *Vibrio campbellii* ATCC BAA-1116 (BB120) genome reference sequence using bowtie2 version 2.4.2 (95). More than 93% of the cleaned reads aligned to the reference sequence. Differential Expression Analysis was performed using DESeq2 ver. 1.44.0 (96). Read pairs mapping concordantly and uniquely to the annotated genes were counted using featureCounts tool ver. 2.0.0 of subread package (97) which was used as input for DESeq2. The following six comparisons were performed: 1) Δ*cap* Δ*luxR* vs Δ*cap,* 2) Δ*cap* Δ*luxR* vs Δ*luxR,* 3) Δ*cap* Δ*luxR* vs WT, 4) Δ*cap* vs WT, 5) Δ*luxR* vs Δ*cap,* and 6) Δ*luxR* vs WT. The comparisons naming convention was treatment vs control. Fold changes for any given gene were computed: as normalized read count in Treatment / normalized read count in Control. Each set of triplicates formed distinct clusters during a principal component analysis, indicating the replicates within each sample being consistent.

Dataset S1 and S2 are excel files that contain the following DESeq2 output for each comparison showing genes that were significantly differentially expressed at 5% FDR: gene ID, gene annotation information, fold-change, Log2 fold-change, *p*-value, adjusted *p*-value: Benjamini Hochberg method (FDR), adjusted p-value - Independent Hypothesis Weighting (IHW) (98), normalized read counts for each individual replicate, normalized read counts for each sample averaged across the 3 individual replicates. Dataset S1 contains the genes that met the 4-fold cutoff for regulation for comparisons 1, 2, 4, and 6 and Dataset S2 contains all significant hits for all comparisons.

#### Overnight Bioluminescence Assays and Growth Curves

Overnight cultures were grown in LM and then pelleted for 3 min at 10,000 rpm at room temperature (RT) to wash cells. Cells were then back-diluted 1:10,000 into 200 µL LM or M9 minimal medium with 20 mM glucose in a black-welled clear-bottom 96-well plate. Selective antibiotics, IPTG, and/or cyclic-AMP were added when applicable and the concentrations are documented in figure captions. At least one space was left between samples to avoid light carryover. OD_600_ and bioluminescence were measured every 30 min for 24 hours using either the BioTek Cytation 3 Plate Reader or the BioTek Synergy H1 Plate Reader. The measurements were recorded from a height of 7 mm, with a gain of 135 (for bioluminescence), the lid on, shaking, at 30°C. Two technical replicates were included on every plate and averaged together to give one biological replicate reading.

#### CAP Protein Purification

To purify the protein CAP for western blots, overnight cultures of *E. coli* BL21(DE3) cells containing pAB105 with 6xHis-tagged CAP were diluted 1:100 into 2x 1 L fresh LB with kanamycin (Kan40) medium and grown shaking at 30°C until the OD_600_ reached 0.4-0.6. Expression of His-CAP was induced with 1 mM IPTG, and this was grown shaking at 30°C for 3 hours. Nickel affinity purification and size exclusion chromatography were performed similarly to the published protocol for the purification of LuxR, HapR, and SmcR (99). Peak fractions were collected and pooled. The fractions were dialyzed for 2 hours at 4°C in dialysis buffer (10 mM Tris pH 7.5, 200 mM KCl, 10 mM MgCl2, 10% volume/volume glycerol) with 10 kDa MWCO SnakeSkinTM dialysis tubing. The protein concentration was calculated using the estimated extinction coefficient of the dimer at 37,000 M-1cm-1. Purified CAP was flash frozen in liquid nitrogen and stored at −80°C.

#### LuxR Protein Purification

His-SmcR (LuxR homolog from *V. vulnificus*; 93% amino acid identity, (10)), was purified as previously described (99) by overexpressing in BL21(DE3) cells containing pJN08 (His-SmcR).

#### Western blots

Whole cell lysates were collected from cells grown to an OD_600_ of 1.0 and separated on Novex 12% Tris-Glycine protein gels (Invitrogen). Proteins were transferred to 0.45 µM nitrocellulose membranes via wet transfer in transfer buffer (48 mM Tris Base, 39 mM Glycine, 0.037% SDS, 20% methanol) at 10V for 65 min. Membranes were blocked overnight at 4°C in a 5% milk/1x TBS-T solution (5 g nonfat dry milk per 100 ml buffer, 25 mM Tris-Cl pH 8.0, 125 mM NaCl, 0.1% Tween 20). Membranes were washed 3x 5 min rocking in 1x TBS-T at RT and then treated with a 1:10,000 dilution of monoclonal anti-RpoB (BioLegend) as a loading control, a 1:3,000 dilution of polyclonal anti-LuxR (71), or a 1:10,000 dilution of monoclonal anti-CRP (BioLegend) for 1 hour rocking at RT. After washing the membranes for 3x 5 min rocking in 1xTBS-T at RT, the membranes were treated with either a 1:10,000 dilution of IRDye 800CW Goat anti-mouse IgG secondary antibody (RpoB and CAP; LICORbio) or a 1:10,000 dilution of IRDye 800CW Goat anti-rabbit IgG secondary antibody (LuxR; LICORbio) for 30 min rocking at RT in dark boxes. All primary and secondary antibody dilutions were dissolved in 1x TBS-T. The membranes were washed 3x 5 min rocking at RT in dark boxes. The membranes were directly added to the LI-COR Odyssey M Imaging System 3340 for visualization and analyzed using LI-COR Empiria Studio 3.0 software.

#### Endpoint Fluorescence Assays

Overnight bacterial cultures were pelleted for 3 min at 10,000 rpm at RT and were washed twice and resuspended in equal volume 1x saline PBS. Resuspended samples were pipetted (200 µL) into a black-welled clear-bottom 96-well plate, leaving at least two spaces in between samples to avoid light carryover. OD_600_ and GFP measurements were recorded on either the BioTek Cytation 3 Plate Reader or the BioTek Synergy H1 Plate Reader from a height of 7 mm using autogain. Excitation and emission for GFP was set to 485 and 528, respectively. Two technical replicates were included on every plate and averaged together to get one biological replicate reading.

#### Chromatin Immunoprecipitation (ChIP) and ChIP-Sequencing

The chromatin immunoprecipitation (ChIP) procedure for *V. campbellii* was modified from a published *Bacillus subtilis* procedure (100) and our previously described *A. fischeri* protocol (101), with minor modifications. Briefly, overnight cultures were back-diluted into 50 mL LM and grown until they reached an OD_600_ of 1.0. Forty-seven mL of cells were crosslinked with formaldehyde at 1% final concentration and rocked for 30 min at RT. The reactions were quenched with 0.5 M glycine and rocked for 5 min at RT. Cultures were pelleted at 4°C, washed twice with cold 1× TBS, frozen with liquid nitrogen, and stored at −80°C until needed. During lysis, pellets were resuspended in ChIP Solution A (12.5 mM Tris pH 8, 12.5 mM EDTA, 62.5 mM NaCl, 25% sucrose, and 1 mM PMSF) with 1 mg/mL lysozyme for 45 min at 37°C. An equal volume of 2× IP Buffer (50 mM Tris pH 7.5, 5 mM EDTA, 150 mM NaCl, 1% Triton) was added to the solution, followed by 1 mM PMSF and 1× Protease Inhibitor Cocktail (Sigma) (49).

Samples were sonicated twice at 4°C using a Qsonica Q800R2 Water Bath Sonicator for 20 min each at 70% amplitude with 10 s on/off pulses. Sonicated lysates were then centrifuged to remove cell debris and pre-cleared by rotating for 1 h at 4°C using 50 µL washed Protein A (for LuxR) or Protein G (for CAP) Magnetic Sepharose beads (Cytiva). Pre-cleared lysates were incubated with either 4 µL anti-LuxR (71) or 4 µL of anti-CRP (BioLegend) and rotated overnight at 4°C. The next day, 50 µL of either Protein A (LuxR) or Protein G (CAP) magnetic beads were added to the lysate-antibody mixtures and rotated for 1 h at 4°C. The resin was washed 3 times with 1× IP Buffer, three times with Wash Buffer (50 mM Tris pH 7.5, 1 mM EDTA, 500 mM NaCl, 1% Triton), once with 1× TE, and eluted with TES (50 mM Tris pH 8, 10 mM EDTA pH 8, 1% SDS) at 65°C for 20 min. Crosslinks were reversed at 65°C overnight and then treated with 0.2 mg/mL RNaseA (NEB) and 0.2 mg/mL Proteinase K (NEB). DNA was extracted using a phenol:chloroform:isoamyl alcohol (25:24:1) mixture with ethanol precipitation, then purified using AmpureXP beads (Beckman Coulter, Inc.). Extracted DNA was then either used for ChIP-seq or ChIP-qPCR. For input samples, 3 mL of cells were collected before crosslinking, and gDNA was extracted using the ThermoFisher GeneJet DNA Purification Kit and diluted to ∼10 ng/μL. The DNA was sonicated in two 6-min rounds at 20% amplitude. Input samples were stored at −20°C until library preparation. For ChIP-seq, Input and IP libraries were prepared for Illumina NextSeq500 or NextSeq2000 high-throughput paired-end sequencing using the NEBNext Ultra II DNA Library Prep Kit for Illumina (NEB #E7645) and sequenced at the Indiana University Center for Genomics and Bioinformatics (CGB). Two biological replicates were sequenced for each sample.

#### ChIP-qPCR

The ChIP-qPCR procedure was modified from a procedure previously described by Chaparian and colleagues (49). Rather than purification using AmpureXP beads, extracted Input and IP DNA was subjected to cleanup using a Qiagen PCR Purification kit. qPCR reactions consisted of a final volume of 20 μL and included 5 ng of DNA sample, 2.5 μM primer mix, 10 μL SensiFAST SYBR Hi-Rox mix (Bioline), and nuclease-free water up to volume. Primers were designed to amplify ∼100bp within the promoters for *cyaA* and *rpoD*. The negative control was selected as *hfq* since CAP does not bind to the *hfq* promoter. All qPCR primers are located in Table S6. qPCR reactions were run in the StepOne Plus RT-PCR Instrument (Applied Biosystems) with the following settings: Step 1: 50°C for 2 min, Step 2: 95°C for 10 min, and Step 3: 95°C for 1 sec, 60°C for 30 sec. Step 3 was repeated for a total of 40 cycles. Fold enrichment was calculated using the ΔΔ*C_T_* method. Three biological replicates were performed with two technical replicates per plate that were averaged together.

#### ChIP-Seq Analysis

ChIP-seq data analysis was conducted using a conda environment and an R (102) (v4.5.3) script. Raw sequencing reads were aligned to the *Vibrio campbellii* ATCC BAA-1116 reference genome utilizing Bowtie2 (95) (v2.5.5). Immunoprecipitated (IP) samples were processed using default end-to-end alignment, while control samples were aligned utilizing the local alignment parameter (--local). The difference in parameters for alignment was chosen to maximize confidence in the called binding sites and minimize false positives. We accept the possibility that biological relevant binding may have been excluded via this technique. Following alignment, reads were filtered utilizing SAMtools (103) (v1.23.1) to retain exclusively properly paired alignments with a mapping quality (MAPQ) score of at least 20 (-f 2 -q 20). Peak calling was executed using MACS2 (104, 105) (v2.2.9.1) in paired-end mode (-f BAMPE), leveraging an empirically calculated genome size, automatic duplicate handling (--keep-dup auto), and a q-value threshold of 0.05. Subsequent peak annotation and replicate consolidation were performed in R relying on the tidyverse (106) (v2.0.0), GenomicRanges (107) (v1.62.1), and rtracklayer (108) (v1.70.1) packages. Narrow peaks exceeding the significance threshold (-log10 q-value ≥ 1.301) were merged across replicates, preserving the peak summit and metadata corresponding to the maximum fold enrichment. The resulting consensus peaks were mapped against GenBank and RefSeq genome annotations to identify overlapping loci and calculate the absolute distance from the peak summit to the nearest flanking start codons (ATGs) on both the upstream and downstream strands. Finally, cross-strain peak overlap events were quantified, and comprehensive summary tables were exported utilizing openxlsx (v4.2.8.1). Input and IP reads were mapped to the *V. campbellii* ATCC BAA-1116 reference genome using JBrowse2 Version 3.6.4 (109) for visualization.

#### Direct Regulon Analysis

Multi-omics integration of RNA-seq and ChIP-seq datasets to define Cap and LuxR direct regulons was executed using a custom pipeline in R (102) (v4.5.3) utilizing the DESeq2 (96) (v1.50.2), tidyverse (106) (v2.0.0), and ashr (110) (v2.2-63) packages. Raw RNA-seq count matrices were imported and subjected to differential expression analysis via DESeq2 (96). To accurately evaluate effect sizes across single and combinatorial knockouts, log2 fold changes (LFC) for five distinct contrasts (Δ*cap* vs. WT, Δ*luxR* vs. WT, Δ*cap* Δ*luxR* vs. WT, Δ*cap* Δ*luxR* vs. Δ*cap*, and Δ*cap* Δ*luxR* vs. Δ*luxR*) were subjected to adaptive shrinkage using the ashr estimator. Genes were classified as differentially expressed utilizing a strict false discovery rate (FDR) threshold of < 0.05 and an absolute shrunken LFC > 1.0. To robustly identify redundantly regulated targets, a positive “null” differential response was explicitly defined as having an FDR ≥ 0.05, an absolute shrunken LFC < 0.5, and a minimum baseMean expression of 20 to ensure adequate statistical power. Concurrently, specificity-filtered ChIP-seq peaks—retained exclusively if absent in their cognate deletion control backgrounds—were mapped to target loci via direct overlap or proximity to the nearest flanking start codons (ATGs). Finally, a combinatorial classification algorithm integrated the binary binding status of CAP and LuxR with the specific expression profiles across all five contrasts. This facilitated the assignment of each locus into discrete regulatory modes, distinguishing between direct additive, cooperative, redundant, antagonistic, and factor-dominant regulatory interactions, while actively controlling for indirect cross-regulation propagated between the transcription factors themselves. Complete diagnostic classifications and summary metrics were subsequently exported utilizing the writexl (v1.5.4) package.

#### Statistical Analyses

Data were plotted and analyzed using GraphPad Prism Software. Specific tests to calculate area under the curve and test for normal distribution and significance are indicated in the figure legends and were performed using GraphPad Prism version 10.6.1.

## Data Availability

RNA-seq and ChIP-seq data were deposited at the National Center for Biotechnology Information (NCBI) Gene Expression Omnibus (GEO) (111). RNA-seq data is under accession GSE339401 and ChIP-seq data is under accession GSE341289. Both datasets can be accessed under the super series accession number GSE341290.

## Author Contributions

**Chase M. Mullins,** conceptualization, formal analysis, investigation, methodology, validation, visualization, writing - original draft, writing - review and editing. **Alyssa S. Ball,** conceptualization, formal analysis, investigation, methodology, validation, visualization, writing - original draft, writing - review and editing. **Logan J. Geyman,** data curation, formal analysis, investigation, methodology, software, validation, visualization, writing-review and editing. **Lydia K. Lukich,** investigation, methodology, validation, visualization, writing - original draft, writing - review and editing. **Lydia K. Hermann,** investigation, methodology, validation, visualization, writing - review and editing. **Ram Podicheti,** data curation, formal analysis, investigation, methodology, resources, software, validation, visualization, writing - original draft, writing - review and editing. **Zhongqing Ren**, investigation, methodology, resources, validation, visualization, writing - review and editing. **Xindan Wang**, conceptualization, funding acquisition, methodology, resources, software, supervision, validation, writing - review and editing. **Douglas B. Rusch,** data curation, formal analysis, investigation, methodology, project administration, resources, software, supervision, validation, visualization, writing - original draft, writing - review and editing. **Julia C. van Kessel,** conceptualization, formal analysis, funding acquisition, methodology, project administration, resources, supervision, validation, visualization, writing - original draft, writing - review and editing.

## BioRender Publication License Acknowledgement

“Figure 8. Model of CAP and LuxR co-regulation of P*_lux_*.” created in BioRender. (https://BioRender.com/op4mmsg) is licensed under CC BY 4.0.

## Acknowledgements

Thank you to Zach Celentano for assisting with protein purification. Thank you to members of the IU Center for Genomics and Bioinformatics for library preparation and sequencing. Thank you to everyone in the collaborative IU Micro community who have provided valuable guidance for this project, with a special thank you to all members of the van Kessel Lab, James McKinlay, Daniel Kearns, Clay Fuqua, and Ankur Dalia. Thank you to summer 2026 IU Writing Tutorial Services (WTS) article writing group members for providing valuable feedback on this manuscript.

