## Supplemental Information for "Catabolite Activator Protein and quorum sensing cross-control group behaviors in *Vibrio campbellii*"

Supplemental Figures (Figures S1-S7)
Supplemental Tables (Table S1-S7)
Legends for Supplemental Datasets S1-S4
Additional References

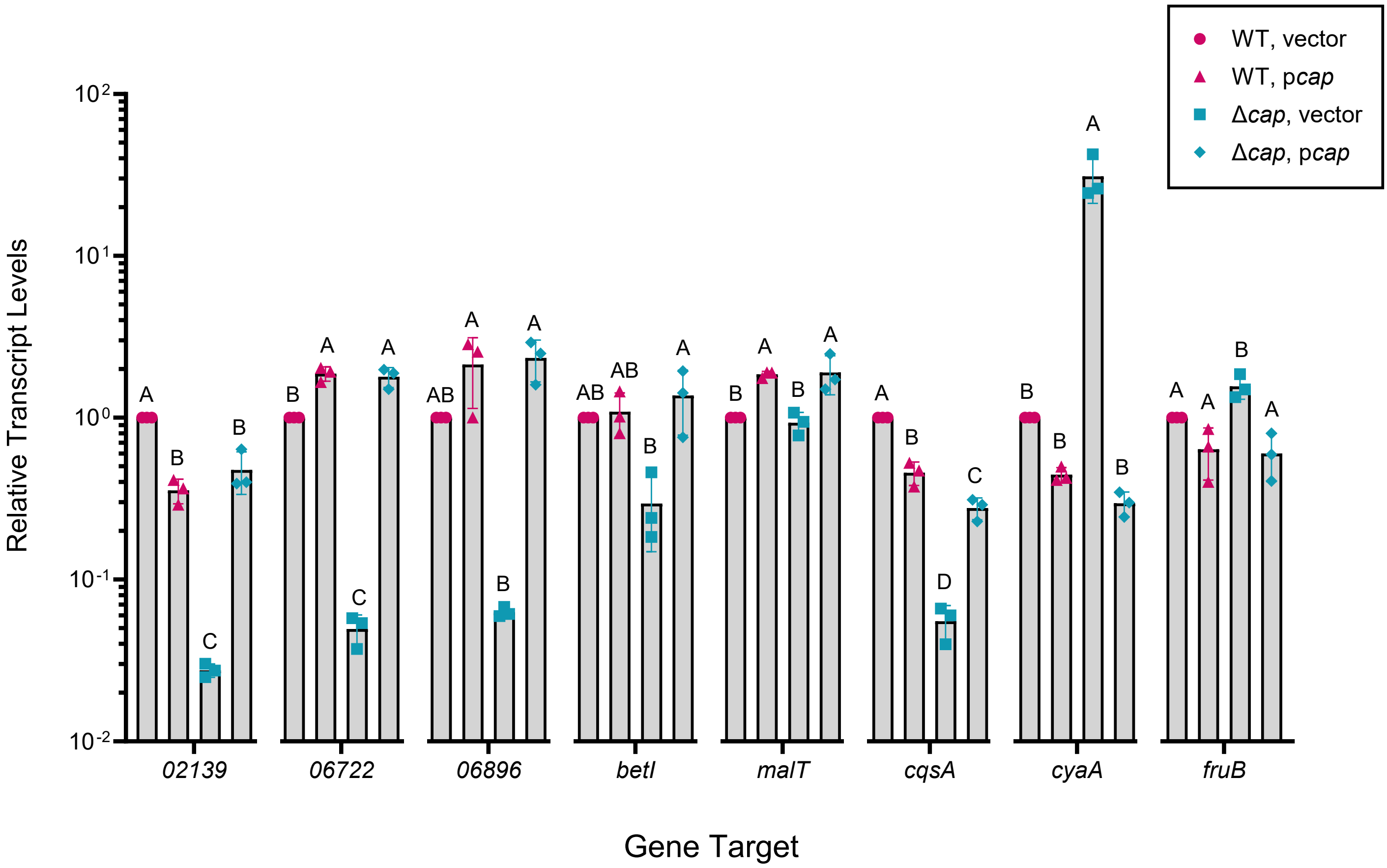

**Supplemental Figure 1.** RT-qPCR analysis of RNA collected at an $OD_{600}$ of 1.0 from WT and Δ*cap V. campbellii* strains harboring either an empty vector (pCS027) or a plasmid with an IPTG-inducible (25 µM) copy of *cap* (pAB100). Relative transcript levels were measured for *VIBHAR_02139, VIBHAR_06722, VIBHAR_06896, betI, malT, cqsA, cyaA,* and *fruB* using the $\Delta\Delta C_T$ method and *hfq* as an internal control. Three biological replicates were performed with two technical replicates averaged together per biological replicate. A one-way ANOVA test was performed on normally distributed data (Shapiro-Wilk test), followed by Tukey's multiple comparisons test between strains within one gene target. Different letters above bars indicate differences in significance between only the strains in each gene target ($p$<0.05; $n$=3). Error bars represent the standard deviation.

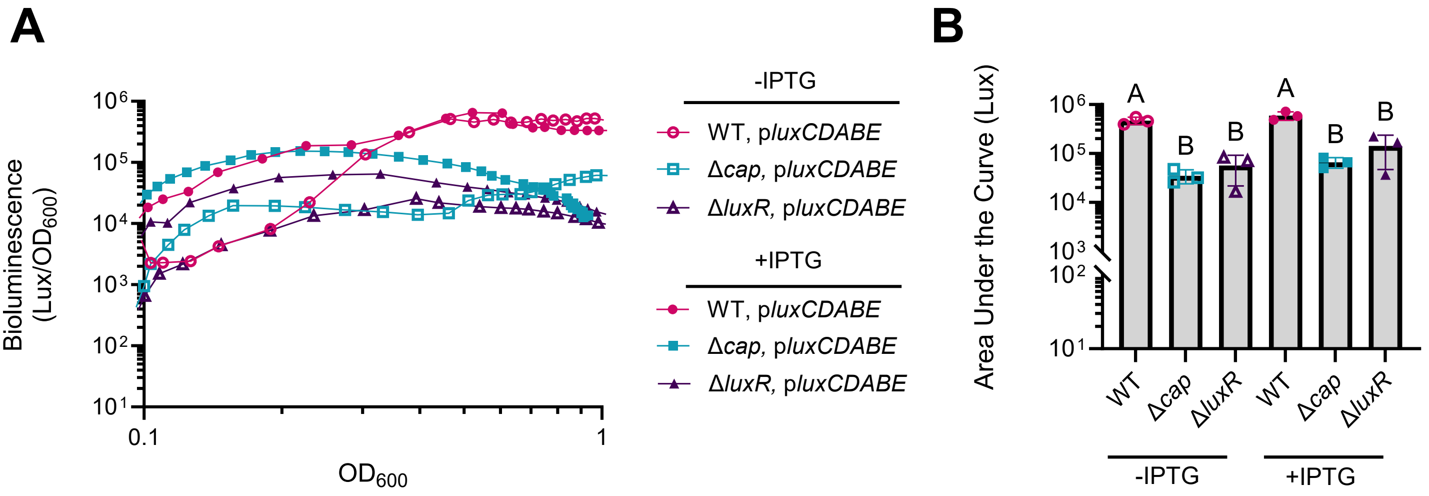

**Supplemental Figure 2.** (A) Bioluminescence production and $OD_{600}$ were measured over 24 hours for Wild-type (WT), Δ*luxR*, and Δ*cap V. campbellii* strains harboring an episomal plasmid expressing *luxCDABE* from an IPTG-inducible promoter (25 μM; pCM024). The graph shown is a representative image from three biological replicates and each line on the graph is an average of two technical replicates. (B) Area under the curve (Lux) was measured from the bioluminescence vs $OD_{600}$ plots in (A) for three biological replicates. A one-way ANOVA test was performed on normally distributed data (Shapiro-Wilk test), followed by Tukey's multiple comparisons test. Different letters above bars indicate differences in significance between strains in pairwise comparisons ($p<0.05$; $n=3$). Error bars represent the standard deviation.

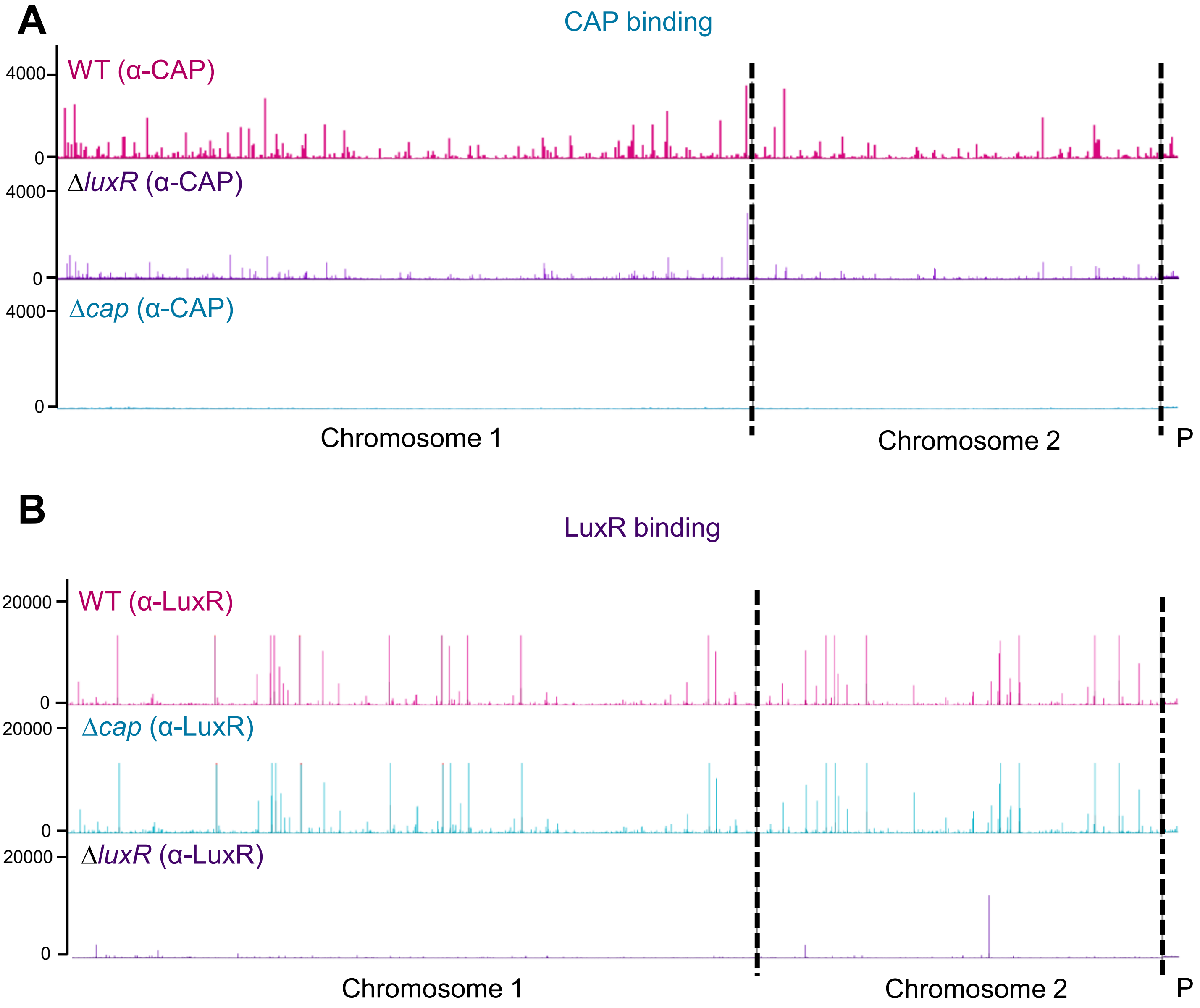

**Supplemental Figure 3.** (A-B) ChIP-seq was performed in WT, Δ*cap*, and Δl*uxR V. campbellii* strains with either α-CAP (A) or α-LuxR antibodies at an $OD_{600}$ of 1.0 (B). Sequenced DNA was mapped to the *V. campbellii* BB120 (ATCC BAA-1116) chromosome and is graphed as enrichment over chromosomal location. The chromosomes are segmented by the black dashed lines. In addition to two chromosomes, *V. campbellii* contains a plasmid which is donated with a P. The Y-axis represents peak height. Screenshots of ChIP peaks were captured using JBrowse 2.

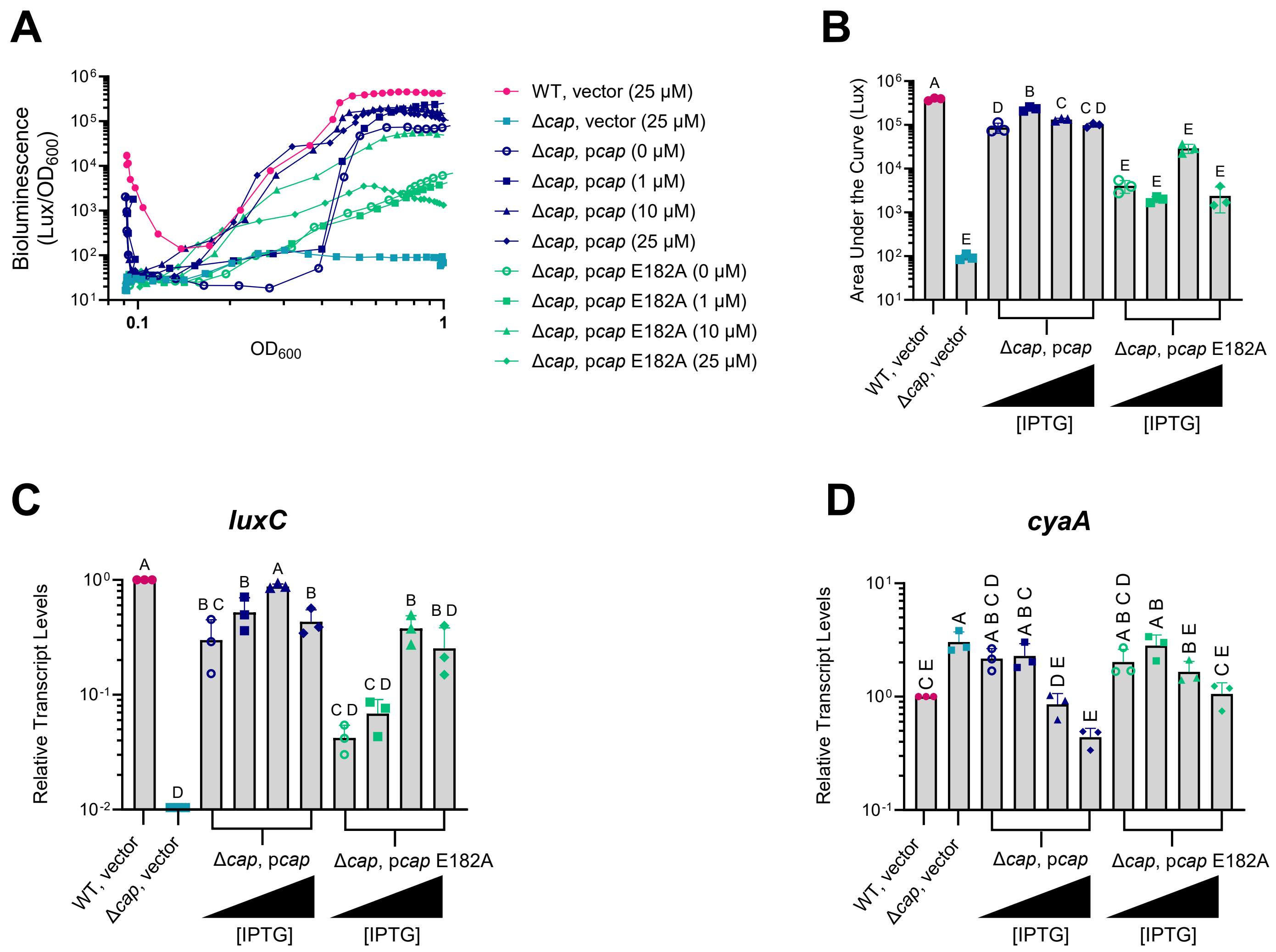

**Supplemental Figure 4.** (A) Bioluminescence production and $OD_{600}$ were measured over 24 hours for Wild-type (WT) and Δ*cap V. campbellii* strains harboring either an empty vector (pCS027) or an episomal plasmid expressing *cap* (pAB100) or *cap* E182A (pCM026) from an IPTG-inducible promoter. IPTG was titrated to final concentrations of 0, 1, 10, or 25 µM as indicated on the graph. The figure is a representative graph from three biological replicates and each line on the graph is an average of two technical replicates. (B) Area under the curve (Lux) was measured from bioluminescence vs $OD_{600}$ plots shown in panel A for three biological replicates. (C, D) RT-qPCR analysis of RNA collected at an $OD_{600}$ of 1.0 from the same strains and conditions as panel A. Three biological replicates were performed with two technical replicates averaged together per each biological replicate. Relative transcript levels were measured for *luxC* (C) and *cyaA* (D) using the $\Delta\Delta C_T$ method and *hfq* as an internal control. (B-D) A one-way ANOVA test was performed on normally distributed data (D'Agostino-Pearson test on the residuals), followed by Tukey's multiple comparisons test between strains. Different letters above bars indicate differences in significance between strains in pairwise comparisons ($p$<0.05; $n$=3). Error bars represent the standard deviation.

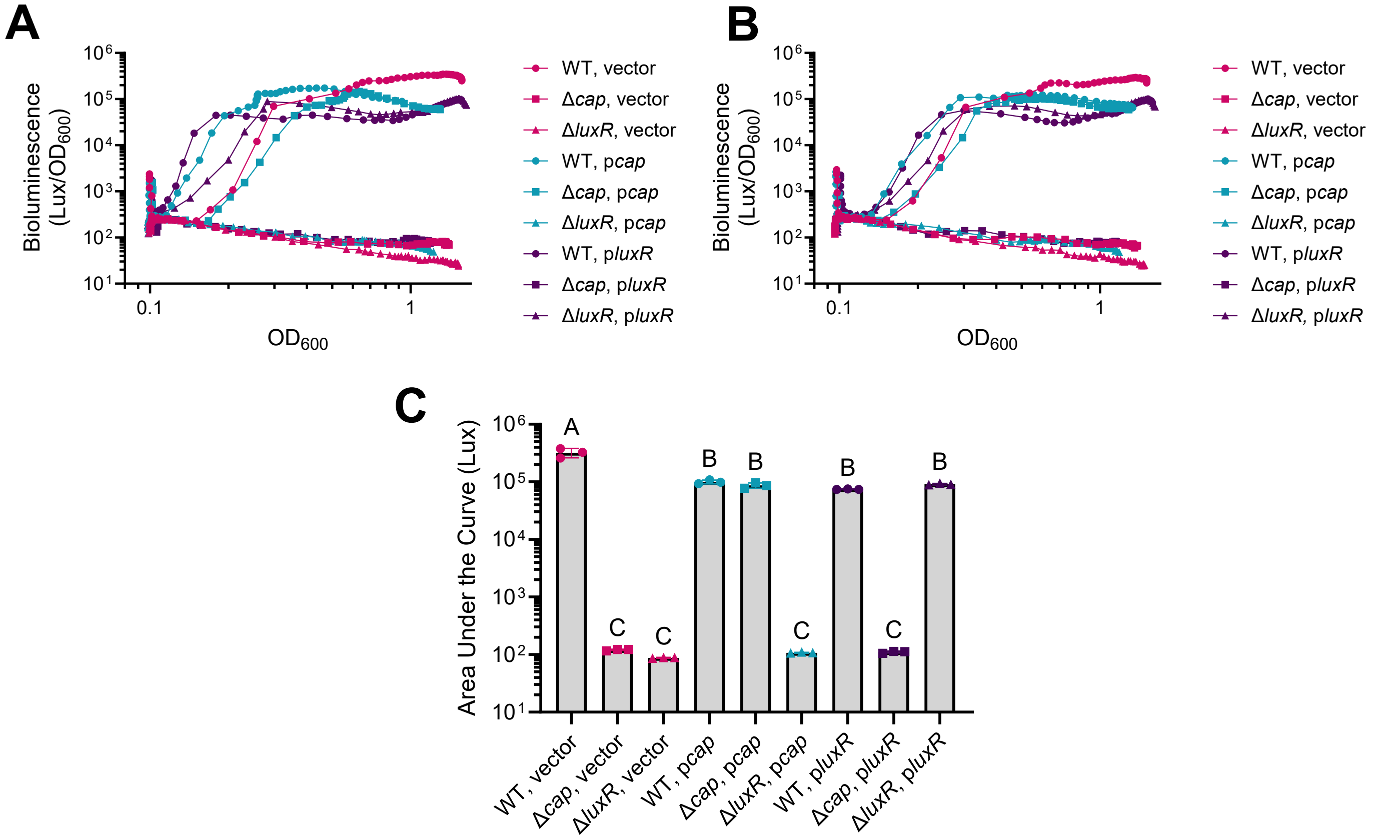

**Supplemental Figure 5.** (A, B) Bioluminescence production and $OD_{600}$ were measured for 24 hours for Wild-type (WT), Δ*luxR*, and Δ*cap V. campbellii* strains harboring either an empty vector (pCS027) or an episomal plasmid expressing *cap* (pAB100) or *luxR* (pJV388) from an IPTG inducible promoter (25 μM). Each line on the graph is an average of two technical replicates. A representative from three biological replicates was presented in the main text as figure 2A; panels A and B constitute the two remaining biological replicates. (C) Area under the curve (Lux) was measured from the bioluminescence vs $OD_{600}$ plots from the three biological replicates presented in panels A-B and Figure 2A. A one-way ANOVA test was performed on normally distributed data (Shapiro-Wilk test), followed by Tukey's multiple comparisons test. Different letters above bars indicate differences in significance between strains in pairwise comparisons ($p<0.05$; $n=3$). Error bars represent the standard deviation.

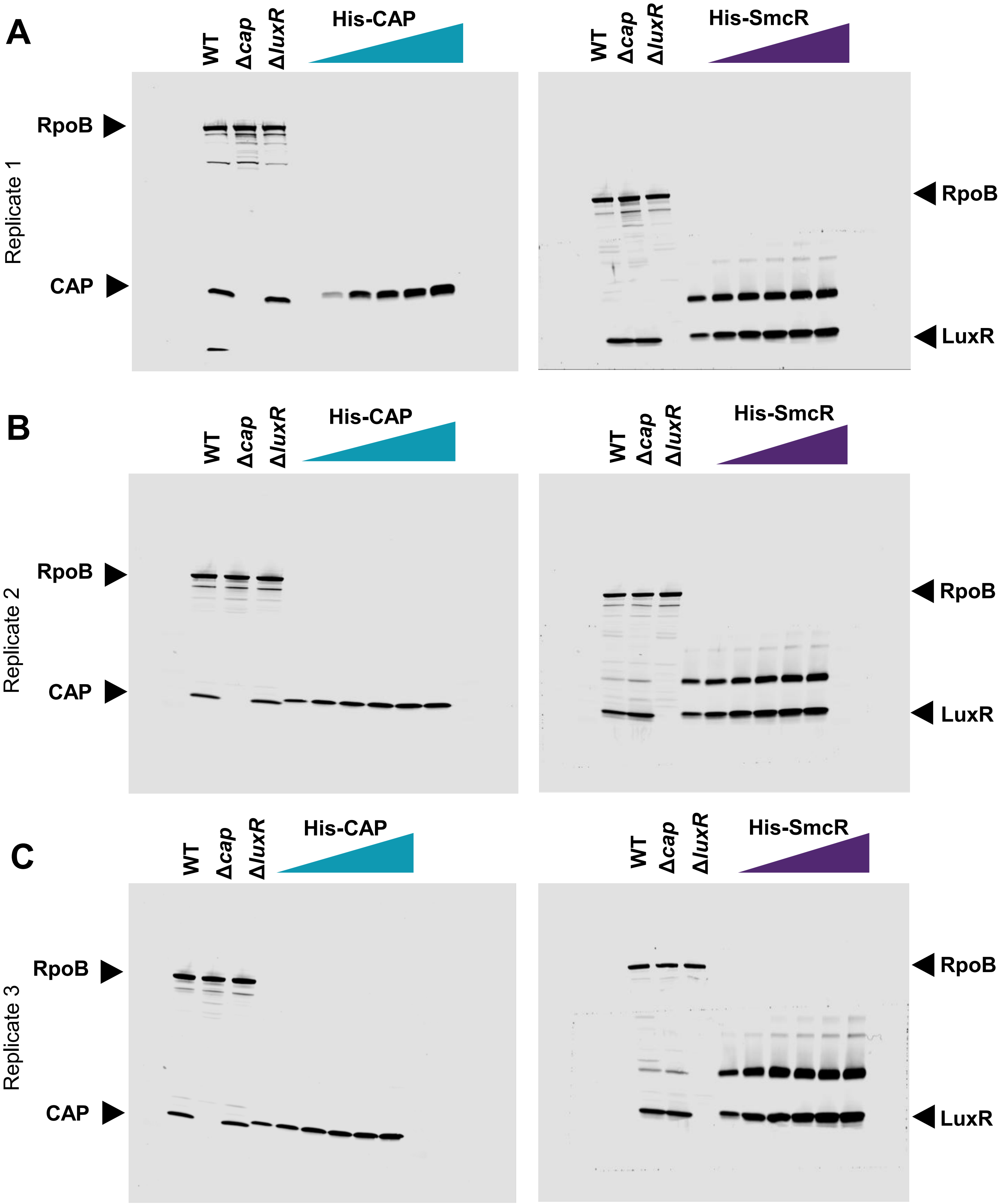

**Supplemental Figure 6.** (A-C) Full membrane images from a western blot analysis of WT, Δ*cap*, and Δ*luxR* lysates collected from cells grown to an $OD_{600}$ of 1.0. A representative, cropped image was presented in figures 2E and 2F in the main text. Lysates were probed using either CAP monoclonal antibodies (α-CAP; left panels) or LuxR polyclonal antibodies (α-LuxR; right panels). RNA polymerase subunit B (α-RpoB) monoclonal antibodies were used to detect RpoB as a loading control in all samples. Increasing concentrations of His-CAP were loaded onto gels to determine a linear range (10 ng, 30 ng, 50 ng, 70 ng, 90 ng, 110 ng for panel A and 20 ng, 40 ng, 60 ng, 80 ng, 100 ng, 120 ng for panels B-C) and are represented by teal triangles. Likewise, increasing concentrations of His-SmcR (the LuxR homolog in *Vibrio vulnificus*; 93% amino acid identity) were loaded onto gels to determine a linear range (50 ng, 100 ng, 150 ng, 200 ng, 250 ng, 300 ng) and are represented by purple triangles. Panel A represents replicate 1 blots, panel B represents replicate 2 blots, and panel C represents replicate 3 blots.

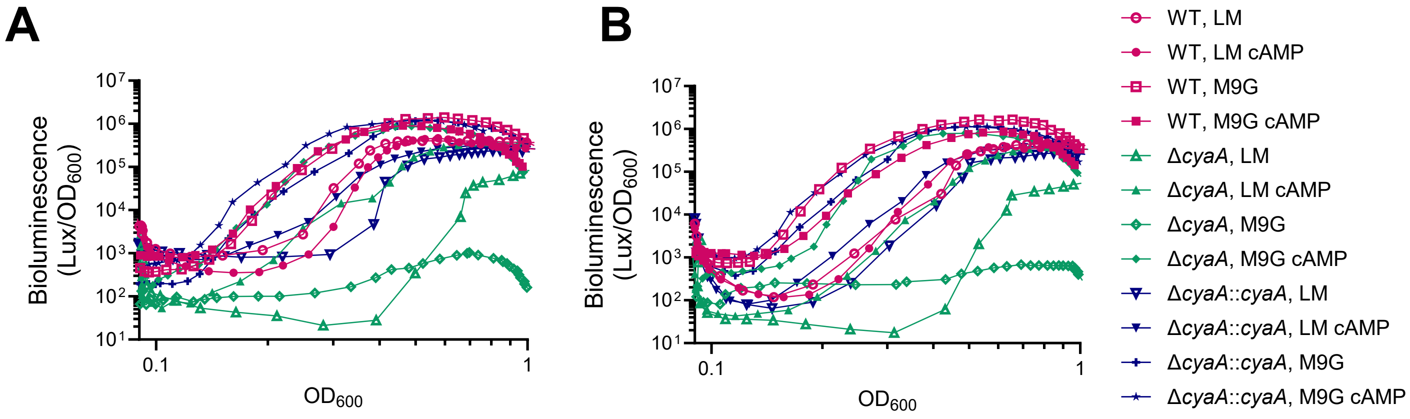

**Supplemental Figure 7.** (A-B) Bioluminescence production and $OD_{600}$ were measured over 24 hours for WT or Δ*cyaA V. campbellii* strains grown in either a rich, undefined medium (LM; LB + NaCl) or M9 minimal medium supplemented with 20 mM glucose (M9G). The Δ*cyaA* strains were complemented with an IPTG-inducible (10 µM) copy of *cyaA* inserted on the chromosome at a non-polar site (Δ*cyaA*::*cyaA*). Exogenous cyclic-AMP (cAMP) was added to the medium at a final concentration of 1 mM. Each line on the graph is the average of two technical replicates. A representative figure was shown in figure 3A in the main text; these are the additional two biological replicates.

**Table S1. MEME Suite-FIMO using *E. coli* CAP consensus sequence identified $P_{luxCDABE}$ CAP Binding Sites.**

| Name | Sequence |
|---|---|
| *E. coli* CAP Consensus | 5'-AAA**TGTGA**TCTAGA**TCACA**TTT-3' |
| CAP Binding site 1 | 5'-AAG**TGCTA**TCTAGG**TGAGA**GGA-3' |
| CAP Binding site 2 | 5'-TAT**AGTGT**TACAAA**TAACA**TTA-3' |
| ($p < 0.05$) | |

a. Bolded nucleotides indicate where the most important CAP-DNA contacts are made as determined by Gartenberg and Crothers 1988 (1) and Benoff *et al.* 2002 (2).

**Table S2. MEME Suite-FIMO using *V. cholerae* CAP consensus sequence identified $P_{luxCDABE}$ CAP Binding Sites.**

| Name | Sequence |
|---|---|
| *V. cholerae* CAP Consensus | 5'-WK**TGA**NNNNNV**TCA**MW-3' |
| CAP Binding site 2 | 5'-AG**TGT**TACAAA**TAA**CA-3' |
| CAP Binding site 3 | 5'-TT**TGA**TCTGCT**TAA**TA-3' |
| ($p < 0.05$) | |

a. Bolded nucleotides indicate where the most well-defined base pairs in the *V. cholerae* CAP consensus sequence as determined by Manneh-Roussel et al., 2018 (3).
b. W= Adenine/Thymine, K= Guanine/Thymine, V=Guanine/Cytosine/Adenine, M= Adenine/Cytosine, N= any DNA nucleotide

**Table S3. MEME Suite-FIMO using *V. campbellii* CAP consensus sequence identified false-positive $P_{luxCDABE}$ CAP Binding Site.**

| Name | Sequence | ***p*-value** | ***q*-value** |
|---|---|---|---|
| *V. campbellii* CAP Consensus | 5’-TAATCGCCGATCAAA-3' | | |
| CAP Binding site 3 | 5'-TTTGATCTGCTTAATA-3' | 0.00261 | 0.76 |
| CAP binding site within *cyaA* promoter (positive control) | 5’-TATTCGCCTATAACT-3’ | 0.000621 | 0.331 |
| CAP binding site within *rpoD* promoter (negative control) | 5’- TATCAGCCGAAGAGA -3’ | 0.00261 | 0.74 |

**Table S4. Strains used in this study.**

| Strains | Genotype | Reference |
|---|---|---|
| ***E. coli* strains** | | |
| S17-1λ*pir* | wild-type | (4) |
| DH10B | wild-type | Life Technologies |
| BL21(DE3) | wild-type | NEB |
| RLG849 | MG1655 wild-type | Gourse Lab unpublished |
| RLG4771 | MG1655 Δ*cap* $Cm^R$ | (5) |
| ***V. campbellii* strains** | | |
| BB120 | wild-type | (6) |
| KM669 | Δ*luxR* | (7) |
| AB003 | Δ*cap* | This study |
| CM056 | Δ*cap* Δ*luxR* | This study |
| CM058 | Δ*cyaA* | This study |
| CM075 | Δ*cyaA*::$P_{IPTG}$-*cyaA* | This study |

**Table S5. Plasmids used in this study.**

| Name | Description | Reference |
|---|---|---|
| pMMB67EH | $P_{tac}$ promoter empty vector, IncQ origin, $kan^R$ | (8) |
| pJV021 | $P_{tac}$ promoter vector, p15a origin, $kan^R$ | (9) |
| pRE112 | gene deletion vector, $cm^R$, *sacB* (sucrose intolerant), R6Kg origin | (10) |
| pJMP1339 | dCas9, R6Kg origin, $kan^R$ | (11) |
| pJMP1039 | Tn7 Transposase, R6Kg origin, $amp^R$ | (11) |
| pCS027 | Empty vector pMMB67EH, $kan^R$ | (12) |
| pAB100 | *V. campbellii* $P_{tac}$-*cap* in pMMB67EH, $kan^R$ | This study |
| pJV388 | *V. campbellii* $P_{tac}$-*luxR* in pMMB67EH, $kan^R$ | (13) |
| pAB105 | *V. campbellii* $P_{tac}$-6xHis-CAP in pET28b, $kan^R$ | This study |
| pCM022 | $P_{tac}$-*cyaA* in pJMP1339, $kan^R$ | This study |
| pCM024 | $P_{tac}$-*luxCDABE* in pMMB67EH, $kan^R$ | This study |
| pCS019 | $P_{luxC}$-*gfp* in pMMB67EH, $kan^R$ | (12) |
| pAB101 | $P_{-216luxC}$-*gfp* in pMMB67EH, $kan^R$ | This study |
| pAB102 | $P_{-156luxC}$-*gfp* in pMMB67EH, $kan^R$ | This study |
| pCS042 | $P_{luxC}$-*gfp* in pMMB67EH, $Gent^R$ | This study |

| | | |
|---|---|---|
| pAB118 | $P_{-216luxC}$-GFP in pMMB67EH, *Gent*$^R$ | This study |
| pAB119 | $P_{-156luxC}$-GFP in pMMB67EH, *Gent*$^R$ | This study |
| pLAFR2 | Empty vector cosmid, *tet*$^R$ | (7, 14) |
| pKM699 | $P_{luxR}$-*luxR* (2.3 kbp fragment) in p*LAFR2* cosmid*, tet*$^R$ | (7, 15) |
| pCM026 | *V. campbellii* $P_{tac}$-*cap* E182A in pMMB67EH, *kan*$^R$ | This study |

**Table S6. RT-qPCR and qPCR primers used in this study.**

| Name | Sequence | Notes |
|---|---|---|
| JCV737 | TGGTTAACAGTGTTCTTCAATAGGATC | *hfq* reverse |
| JCV738 | ATGGCTAAGGGGCAATCTC | *hfq* forward |
| CM128 | AACACGGCAACATTACTCATG | *VIBHAR_02139* forward |
| CM129 | CGAAACAACGTCACACCAAG | *VIBHAR_02139* reverse |
| CM126 | TGCACTTATCGAAGACGACGA | *VIBHAR_06722* forward |
| CM127 | CTAAGAATGGCGGCTTTGCC | *VIBHAR_06722* reverse |
| CM124 | CTGCAACGTATTTACTTGCTG | *VIBHAR_06896* forward |
| CM125 | GGTTGATAGAGGAATCCCCA | *VIBHAR_06896* reverse |
| CM122 | GGTTGGGATGCCTGAAATCA | *betI* forward |
| CM123 | GCTGATCAACGAGATACTCG | *betI* reverse |
| CM120 | GGATTCCATCAAAACTTACTCG | *malT* forward |
| CM121 | GTACCAGTTTGTAGTAAGGCG | *malT* reverse |
| CM118 | CCACTTCCTTCCTTTGTTGAGG | *cqsA* forward |
| CM119 | TGGTCGCTTCCCTAATACCA | *cqsA* reverse |
| CM104 | GCTAAGTAAGAACGACATTACCC | *fruB* forward |
| CM105 | TTCAACCAGACCTTTGTCTGTC | *fruB* reverse |
| CM114 | GCGATTAGATAACCTGAACCAG | *cyaA* forward |
| CM115 | GTTCAACAAGGCAGGTATCAAG | *cyaA* reverse |
| JCV745 | GCAAAGAGACCTCGTACTAGG | *luxR* forward |
| JCV746 | GCGACGAGCAAACACTTC | *luxR* reverse |
| *luxC* right | TGTTCATTTAACTCAGATGGTGACT | *luxC* right |
| *luxC* left | TTCTTCTTGAATACTCTTCGCTCTT | *luxC* left |
| AB336 | TAAACCTCAAACCGACCCAA | *cap* forward |
| AB337 | ACGATGTAGTACAAAGTCTCTGC | *cap* reverse |
| CM167 | AGTGTGACTGGATTCTCATTTAGAC | F $P_{cyaA}$ ChIP-qPCR primer (-232 bp from start of gene) |
| CM168 | CCACACTAGTCGCAAAAATTGATAG | R $P_{cyaA}$ CHIP-qPCR primer (-136 bp from start of gene) |
| CM169 | GCTGCTAGCTTTGATGCTAGA | F $P_{rpoD}$ ChIP-qPCR primer (-133 bp from start of gene) |
| CM170 | GGAATCAGTATGCGAAATGCA | R $P_{rpoD}$ ChIP-qPCR primer (-30 bp from start of gene) |

**Table S7. Genes mentioned in this study and their associated locus tags.**

| Gene Name | Locus Tag |
|---|---|
| *cap* | *VIBHAR_RS00360* |
| *luxR* | *VIBHAR_RS16180* |
| *hfq* | *VIBHAR_RS00455* |
| *luxC* | *VIBHAR_RS23945* |
| *luxD* | *VIBHAR_RS23940* |
| *luxA* | *VIBHAR_RS23935* |
| *luxB* | *VIBHAR_RS23930* |
| *luxE* | *VIBHAR_RS23925* |
| *VIBHAR_02139* | *VIBHAR_RS10035* |
| *VIBHAR_06722* | *VIBHAR_RS25890* |
| *VIBHAR_06896* | *VIBHAR_RS26600* |
| *betI* | *VIBHAR_RS23655* |
| *malT* | *VIBHAR_RS17930* |
| *cqsA* | *VIBHAR_RS23255* |
| *cyaA* | *VIBHAR_RS01465* |
| *fruB* | *VIBHAR_RS22725* |
| *rpoD* | *VIBHAR_RS04015* |
| *focA* | *VIBHAR_RS08700* |
| *glpT* | *VIBHAR_RS15475* |
| *VIBHAR_06505* | *VIBHAR_RS25000* |
| *malE* | *VIBHAR_RS19540* |
| *malF* | *VIBHAR_RS19545* |
| *malG* | *VIBHAR_RS19550* |
| *cqsS* | *VIBHAR_RS23260* |
| *luxM* | *VIBHAR_RS12950* |

**Legends for Supplemental Datasets:**

Supplemental Dataset S1: Genes ≥ 4-fold regulated by CAP and LuxR
Supplemental Dataset S2: Unfiltered RNA-seq hits with all normalized read counts
Supplemental Dataset S3: ChIP-seq peaks ≥ 2-fold enriched by CAP and LuxR
Supplemental Dataset S4: Genes directly regulated by CAP and LuxR
